# In-Depth Resistome Analysis by Targeted Metagenomics

**DOI:** 10.1101/104224

**Authors:** Val F. Lanza, Fernando Baquero, José Luós Martónez, Ricardo Ramos-Ruóz, Bruno González-Zorn, Antoine Andremont, Antonio Sánchez-Valenzuela, Dusko Ehrlich, Sean Kennedy, Etienne Ruppe, Willem van Schaik, Rob J. Willems, Fernando de la Cruz, Teresa M. Coque

**Author notes:** Current affiliation: Hub Bioinformatique et Biostatistique, C3BI & Biomics Pole, CITECH Institut Pasteur, Centre François Jacob, 28 rue du Docteur Roux, 75015 Paris (Sean Kennedy). Current affiliation: Genomic Research Laboratory, Service of Infectious Diseases, Geneva University Hospitals, rue Gabrielle-Perret-Gentil 4, 1205 Geneva, Switzerland (Etienne Ruppe).

## Abstract

We developed ResCap, a targeted sequence capture platform based on SeqCapEZ technology, to analyse resistomes and other genes related to antimicrobial resistance (heavy metals, biocides and plasmids). ResCap includes probes for 8,667 canonical resistance genes (7,963 antibiotic resistance genes and 704 genes conferring resistance to metals or biocides), plus 2,517 relaxase genes (plasmid markers). Besides, it includes 78.600 genes homologous to the previous ones (47,806 for antibiotics and 30,794 for biocide or metals). ResCap enriched 279-fold the targeted sequences detected by metagenomic shotgun sequencing and improves their identification. Novel bioinformatic approaches allow quantifying “gene abundance” and “gene diversity”. ResCap, the first targeted sequence capture specifically developed to analyse resistomes, enhances the sensitivity and specificity of available metagenomic methods to analyse antibiotic resistance in complex populations, enables the analysis of other genes related to antimicrobial resistance and opens the possibility to accurately study other complex microbial systems.

## INTRODUCTION

Antimicrobial resistance is considered a major Global Health challenge recently included in the political agendas of international bodies such as G8 and IMF^1^. The adoption of measures to face the “antibiotic resistance crisis” ^2^ is impaired by the controversy about “what” is resistance and “how” and “where” should be detected and analysed ^3–45^

Metagenomic methods are increasingly used to analyse the ensemble of genes encoding antibiotic resistance in different microbial ecosystems which has recently been defined as the “resistome”^6–16^. An important hurdle of the available resistome analyses is the low discrimination in the detection of minority populations harbouring resistance genes (often present at concentrations below the detection level of the methods used)^17^ and/or the identification of allelic variants that might confer different resistance phenotypes.

A sensitive and specific identification of antibiotic resistance genes in a metagenome background is required for assessing the associated risks in terms of Public Health 18a, 18b. Such methodological challenge parallels the difficulties of analysing sets of orthologous genes of many individuals for the diagnosis of human multifactorial inherited diseases ^19^. In this case, the use of “capture-based” or “targeted” sequencing strategies, was a cost-effective and high-throughput alternative that overcame the limitations of metagenomic shotgun sequencing (MSS)^20,21^. In-solution targeted capture platforms (TCP) take advantage of Next Generation Sequencing to provide technical improvements over array-based platforms or other genome-partitioning approaches in terms of scalability, cost-effectiveness, and enhanced data quality (lower variance in target coverage, more accurate SNP calling, higher reproducibility and longer assembled contigs)^22^. Currently, TCPs offer a tremendous potential for boosting advances in environmental and ecological studies, particularly involving micro-biodiversity research, which requires the isolation of sequences of interest from a mixture of DNAs of a complex multiplicity of organisms^23^.

Our work reports the development and validation of the first TCP for the analysis of bacterial resistomes, which we designate as ResCap (for Resistome Capture). We show that ResCap results in a significant improvement in sensitivity and specificity over previous metagenomic analysis of antimicrobial resistance. ResCap also allows the analysis of the presence and diversity of genes conferring resistance to other antimicrobials (heavy metals and biocides), which are frequently co-selected with antibiotic resistance genes and also genes from replicons of the mobilome (as plasmids). An *ad-hoc* advanced bioinformatics pipeline, developed in parallel, exploits the capabilities of Rescap comparative metagenomic analysis. The metagenomic approach described here opens the way for a series of applications in the identification, epidemiological surveillance, ecology, and study of evolutionary trajectories of resistance genes.

## RESULTS

### Targeted metagenomics, a tool for high-resolution analysis of resistome

ResCap was designed to establish a standarized framework that would allow performing both quantitative and qualitative analysis of resistomes. Also, to facilitate the analysis of novel genes potentially involved in the resistance to antibiotics, metals, biocide or any combination of them. As a proof of concept, we compared the performance of ResCap with metagenomic shotgun sequencing (MSS) by analysing the resistome in ^17^ fecal samples, 9 from humans and 8 from swine.

ResCap exhibits a target capacity (total amount of targeted sequences) of 88.13 Mb, and includes probes for 78,600 non-redundant genes (81,117 redundant genes), including 7,963 functionally validated antibiotic resistance genes, 704 functionally validated metal & biocide resistance genes, and 2,517 relaxase genes (genes used for plasmid identification and classification)^24^. Besides the 8,667 genes that confer functionally proved resistance to antimicrobials (canonical genes), the platform also includes targets for 78,600 homologous of resistance genes (47,806 for antibiotics and 30,794 for biocide and metals resistance). The criteria used to select the targeted genes are explained in the section Material and Methods.

ResCap performance was compared with MSS in two ways. First, by applying a reference-based approach that maps metagenome reads against specific databases (AbR, Metal & Biocides and Relaxases). Second, by applying a reference-free approach that assembles metagenomic reads and performs a functional annotation. The results of both evaluations are detailed below.

### Reference-based evaluation

This section addresses how the abundance and diversity of resistance genes (ResCap or those already validated) were calculated.

#### ResCap achieves better recovery of target genes than MSS

An average of 1.9×10^7^ paired-reads was obtained from the MSS and ResCap datasets (0.92-3.2×10^7^). The on-target average (the number of reads mapping on the target genes relative to the total read number) against the selected databases (see Material and Methods) was 0.11% (0.07-0.18) for MSS data and 30.26% (20.27–41.83%) for ResCap data, which represents an enrichment of 279-fold (**Table 1**).

**TABLE 1.**

| Sample | Metagenome shotgun sequence MSS |  |  |  |  | ResCap |  |  |  |  | Gain |
| --- | --- | --- | --- | --- | --- | --- | --- | --- | --- | --- | --- |
|  | *Nº Reads* | *AbR* | *BacMet* | *Relaxases* | *Total* | *Nº Reads* | *AbR* | *BacMet* | *Relaxases* | *Total* |  |
| Bichat1 | 14,127,290 | 0,05% | 0,002% | 0,04% | 0,10% | 16,705,789 | 19% | 0,48% | 5,22% | 24,68% | 244,24 |
| Bichat2 | 15,128,135 | 0,05% | 0,028% | 0,04% | 0,12% | 33,589,838 | 12% | 5,56% | 2,98% | 20,27% | 170,81 |
| Bichat3 | 14,488,245 | 0,05% | 0,005% | 0,03% | 0,09% | 17,276,637 | 34% | 2,31% | 5,96% | 41,83% | 480,59 |
| Bichat6 | 17,476,666 | 0,07% | 0,001% | 0,05% | 0,12% | 19,191,320 | 25% | 0,85% | 5,84% | 32,13% | 261,87 |
| Bichat7 | 16,732,926 | 0,07% | 0,002% | 0,05% | 0,13% | 28,530,922 | 27% | 0,33% | 5,99% | 33,60% | 267,77 |
| Bichat9 | 17,058,000 | 0,03% | 0,013% | 0,05% | 0,09% | 18,038,257 | 14% | 10,59% | 7,55% | 31,80% | 336,29 |
| Bichat10 | 15,039,883 | 0,03% | 0,066% | 0,06% | 0,15% | 34,798,281 | 6% | 28,81% | 5,48% | 40,63% | 265,17 |
| Bichat11 | 13,425,077 | 0,03% | 0,091% | 0,06% | 0,18% | 35,901,508 | 5% | 26,73% | 4,72% | 35,98% | 201,67 |
| Bichat13 | 17,903,872 | 0,05% | 0,023% | 0,06% | 0,14% | 26,283,052 | 16% | 13,27% | 7,39% | 36,85% | 270,02 |
| F266 | 19,557,955 | 0,06% | 0,005% | 0,02% | 0,08% | 14,024,345 | 21% | 4,62% | 2,57% | 28,23% | 337,50 |
| PIG20 | 27,375,311 | 0,08% | 0,028% | 0,01% | 0,12% | 22,485,364 | 18% | 16,20% | 1,81% | 36,38% | 298,23 |
| PIG26 | 13,831,057 | 0,07% | 0,005% | 0,02% | 0,10% | 15,756,070 | 19% | 3,88% | 2,73% | 25,93% | 271,20 |
| PIG29 | 18,945,765 | 0,09% | 0,018% | 0,02% | 0,12% | 26,223,850 | 18% | 10,26% | 2,52% | 30,65% | 248,31 |
| PIG31 | 12,778,294 | 0,07% | 0,010% | 0,02% | 0,09% | 18,055,019 | 13% | 5,76% | 2,08% | 20,77% | 219,12 |
| PIG528 | 19,689,471 | 0,06% | 0,003% | 0,02% | 0,08% | 13,864,257 | 21% | 2,70% | 3,11% | 26,83% | 323,23 |
| PIG94 | 15,985,219 | 0,07% | 0,004% | 0,02% | 0,10 | 15,351,408 | 18% | 2,57% | 3,31% | 24,05% | 240,20 |
| PIG96 | 9,290,402 | 0,06% | 0,001% | 0,01% | 0,07% | 12,225,935 | 21% | 1,13% | 1,67% | 23,84% | 320,90 |

The analysis of the gene abundance, expressed in RPKMs (reads per kb per million reads, see Materials and Methods), demonstrates better recovery of genes coding for resistance to antibiotics, heavy metals, biocides and relaxases (plasmid genes) when using ResCap than when using MSS. **Figure 1** represents the RPKMs inferred before (MSS) and after capture(ResCap) for all the samples analysed, while **Figure S1** shows the gain plots for each sample. Most canonical genes (99.3%, 1339/1.348) detected by MSS were also detected with ResCap. Furthermore, almost half of the genes detected by ResCap (42%,975/2323), were not detected by MSS. The linearity of the system was evaluated by using a linear regression model for the genes detected in each paired-sample (MSS vs ResCap). An R2 mean of 0.813 (0.85-0.99) shows a good match between both protocols.

**Figure 1.**
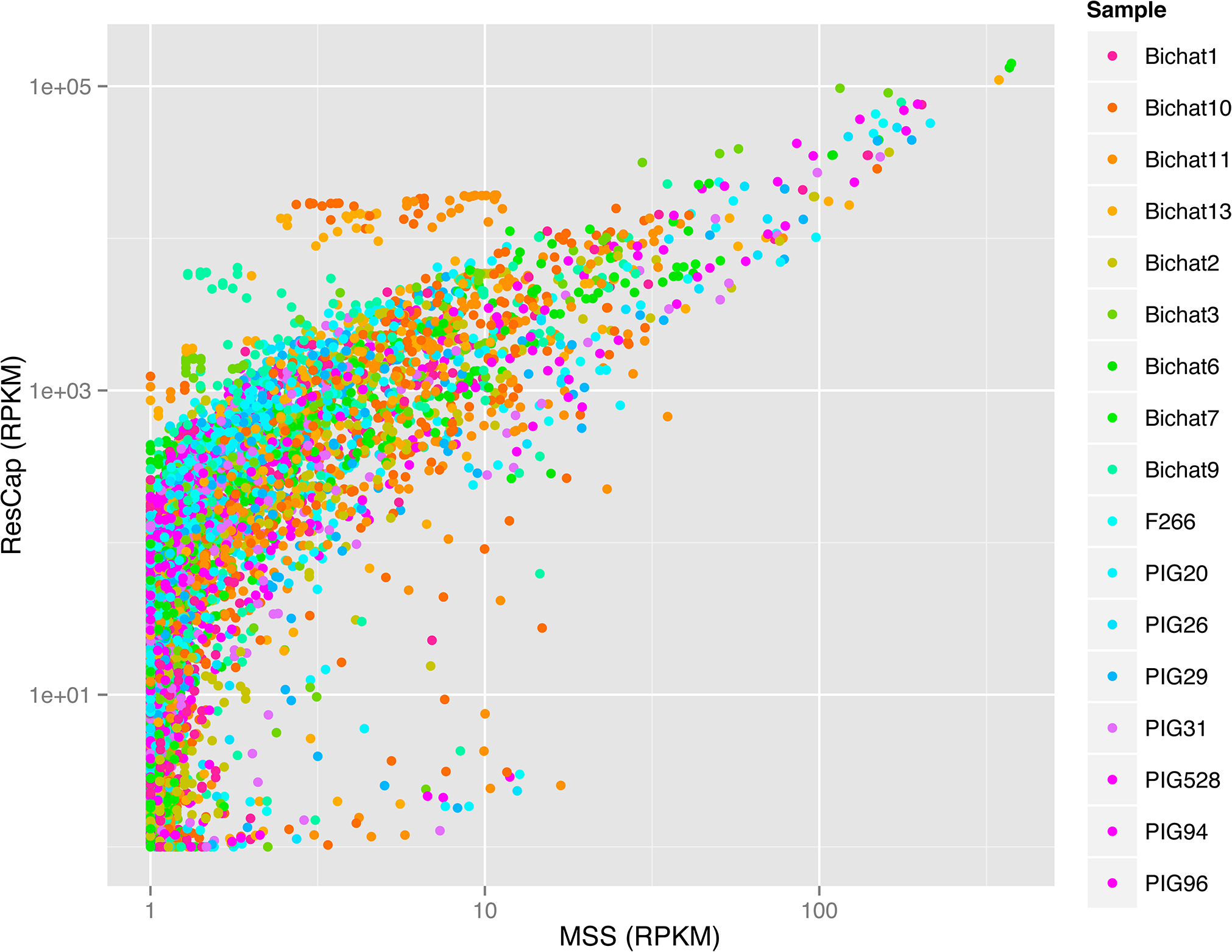
Gain function plot. Representation of the gain in reads per kilobase per million of reads of each detected gene between MSS protocol (abscissa axis) and ResCap (ordinate axis). Genes only detected by ResCap are represented by the dot cluster in the initial values of abscissa axis. The pictures are represented in log-log scale to a better perception of the linearity of the gain function in genes detected by both protocols.

The enrichment of canonical resistance genes when using ResCap was similar in samples from humans and swine. Nonetheless, the differences in the relative abundance of genes encoding resistance to antimicrobials (antibiotics, heavy metals, biocides) and relaxases in different samples (Figure 2) is not surprising due to the variability of microbiotas of different hosts ^25,26^

**Figure 2.**
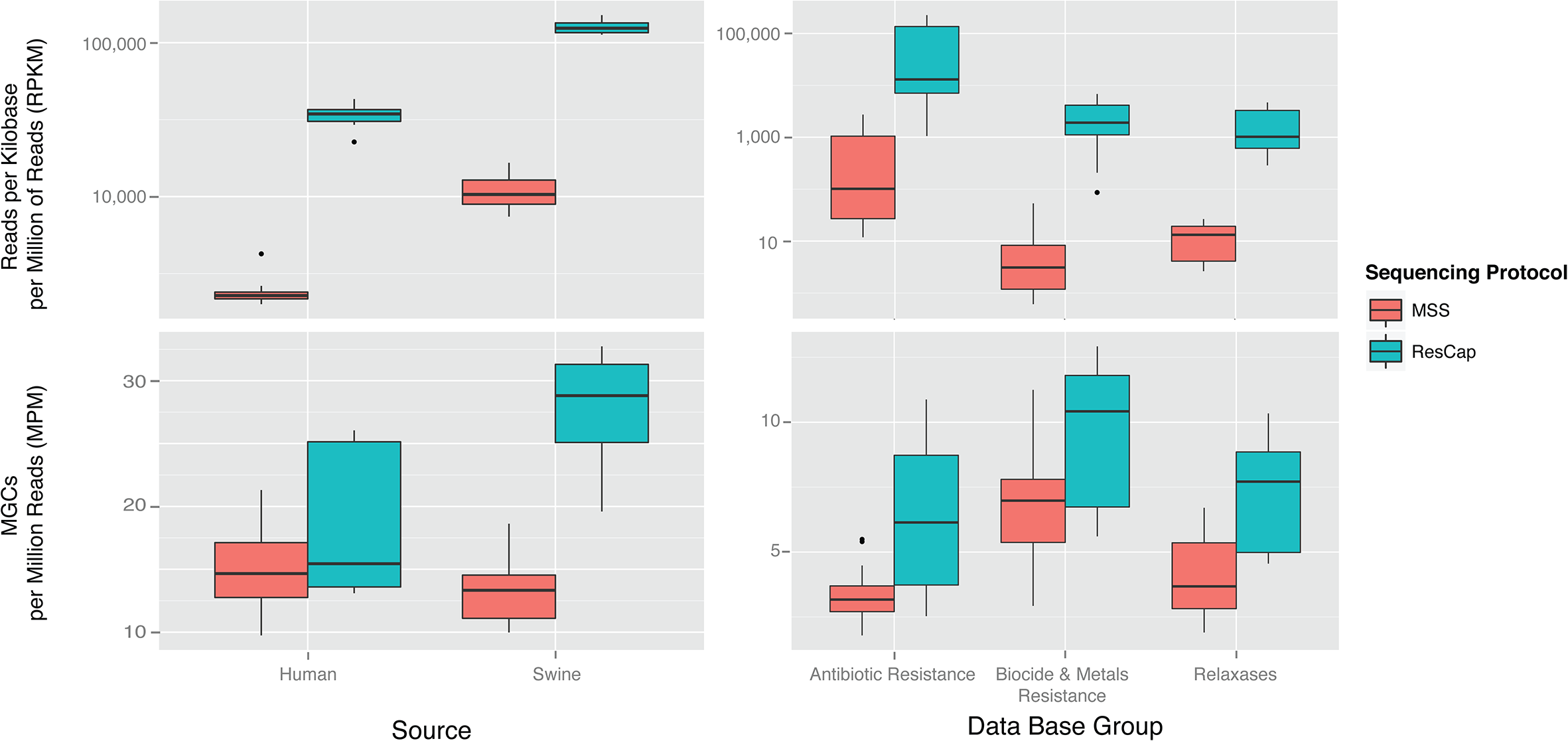
Platform Efficiency by Source Sample and Data Base Group. Data distribution of the platform efficiency evaluating (a) the number of mapped reads per million of sequenced reads against canonical (well-known) genes data set; and (b) the number of detected genes per million of sequenced reads using as reference the well-known genes data set. Fecal samples were differentiated according to the source (9 from humans and 8 from swine). Data distribution of the platform efficiency evaluating (c) the number of mapped reads per million of sequenced reads against the three canonical genes groups and (d) the number of detected genes per million of sequenced reads using as reference the three canonical genes groups.

#### ResCap addresses gene diversity

Allele redundancy of some resistance genes hinders the correct estimation of “gene diversity” and precludes a correct estimation of “gene abundance” in metagenomes when using most available metagenomic tools.

To overcome this issue, we define the term Mapping Gene Cluster (MGC) as the group of alleles/genes detected by the same set of reads (see Material and Methods). MGCs, firstly defined in this work, allow an estimation of gene diversity across samples, and are measured as the number of MGCs per million reads (MPM). The number of MPMs increased 1.3 fold in humans (0.7-1.74) and 2.1-fold (2.3-1.9) in swine when using ResCap instead of MSS (**Figure 2**).

An increase in reads per MGC does not imply a homogeneous distribution of the reads. Therefore, we also determined the “gene horizontal alignment coverage”, which was defined as the fraction of a gene that is covered by reads. The probability of identifying an allele-specific mutation will also increase with the number of reads per nucleotide or “gene depth coverage”. **
Figure 3
** shows the improvement of “gene alignment horizontal coverage” using ResCap and MSS (average= 97.5%, range = 66%-99% vs. average= 73.4%, range = 35.9%-94.8%, respectively). Most genes were almost fully covered by reads and there was also a general increase in “gene depth coverage” (**Figure S2**). As a consequence, the number of genes unequivocally detected by ResCap almost doubled that of MSS (n=26, range 17.1-30.0 genes per sample per million of reads vs. n=14.9, range 12-17.6 genes per sample per million of reads). The number of reads unequivocally mapped increased up to 300 fold (2×105 for ResCap *vs* 8×102 for MSS) (**Figure 4**).

**Figure 3.**
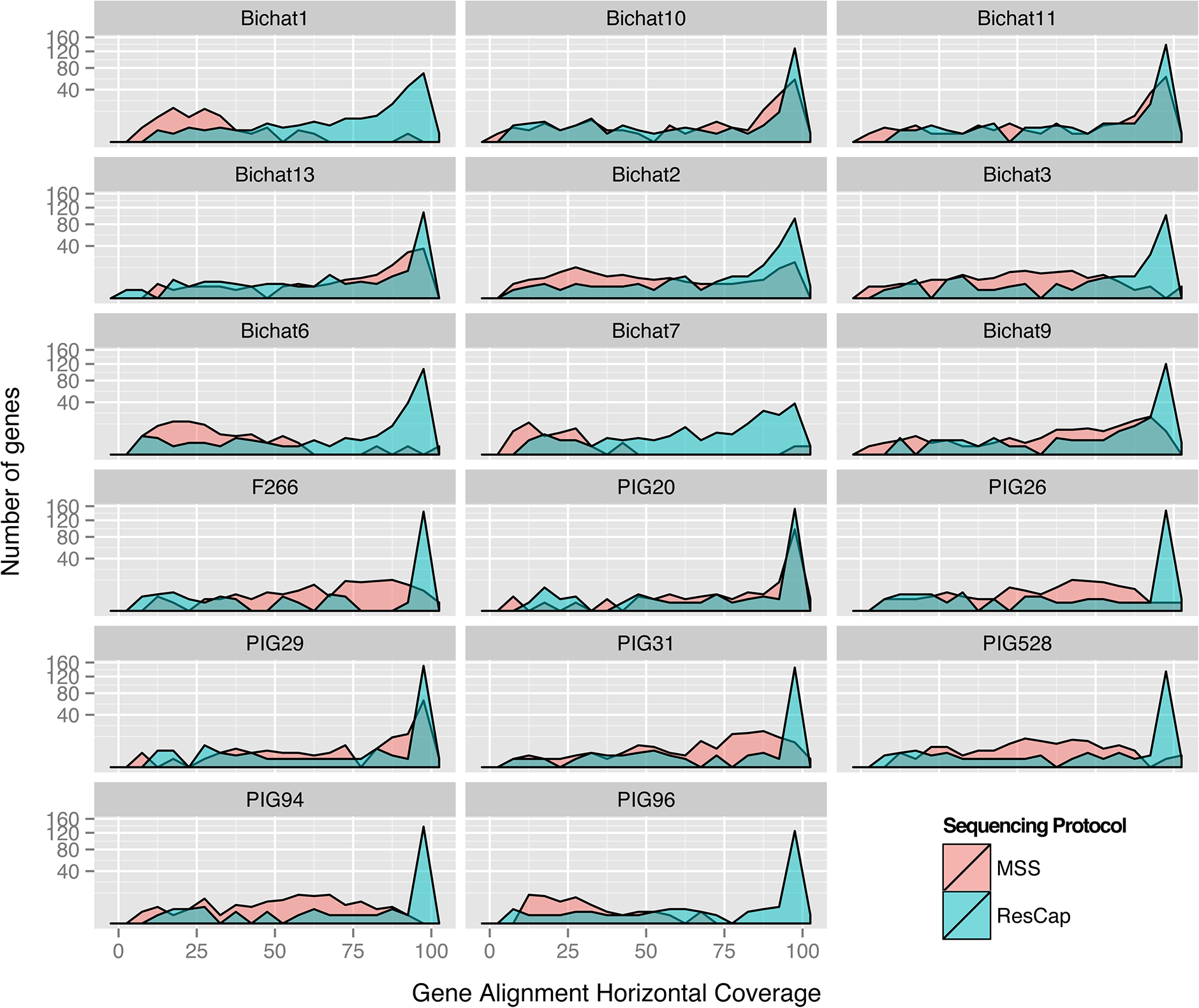
Longitudinal coverage distribution. The figure shows the comparison of longitudinal coverage distribution between protocols in each sample. Distributions are represented by density parameter and expressed by the number of genes (ordinate axis) and coverage percent (abscissa axis).

**Figure 4.**
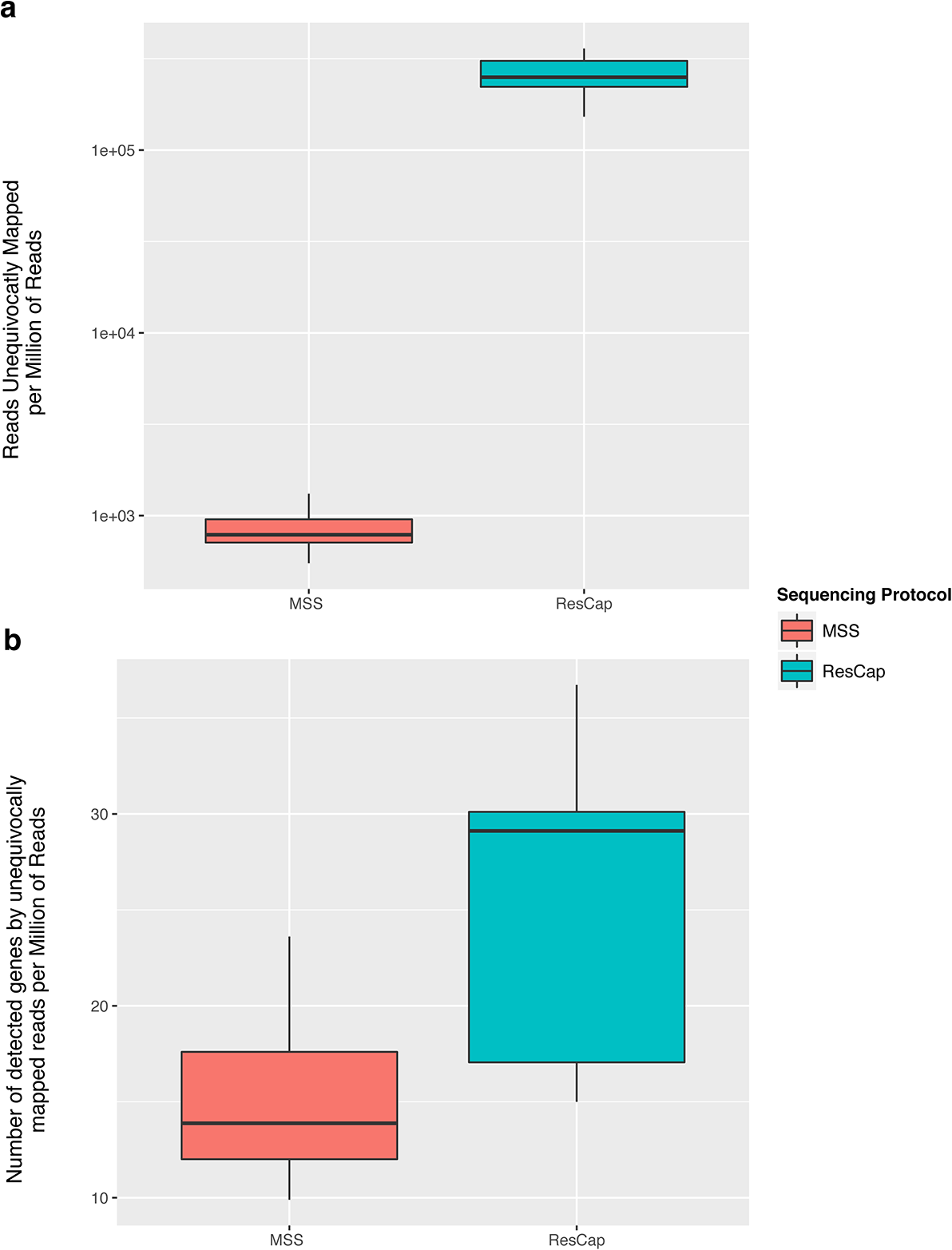
Quantification of unequivocally mapping reads. The figure shows the comparative of the quantification of reads that mapping on just one gene (or allele). First the abundance of reads that are unequivocally mapped on one gene (a). On another hand, the number of genes (or Mapping Gene Cluster) that have almost one read that mapping unequivocally (b). Box plots are differentiated for MSS protocol and ResCap protocol.

**Figures S3** shows the abundance (RPKMs) and diversity (MPMs) obtained by ResCap and MSS for individual categories of resistance genes (antibiotics, biocides and metals), which also illustrates the improved sensitivity of ResCap vs. MSS. **Figure S4** reflects that although both ResCap and MSS can track the most abundant gene families as those conferring resistance to beta-lactams, macrolides, aminoglycosides and tetracyclines, followed by those conferring resistance to phenicols and sulphonamides, many canonical resistance genes were only detected by the ResCap platform (e.g. *mecA, blaZ* in beta-lactams; *ermA, ermC, ermD, erm33* or *lnu* among macrolides; *fexA, catA and catB* alleles among phenicols). Genes encoding resistance to fluoroquinolones, glycopeptides, or trimethoprim, first line antibiotics families used to treat community and hospital-based infections, were barely detected using MSS but unequivocally detected with ResCap (e.g. *dfrA^16^, dfrA^15^, dfrG, dfrK* among those conferring resistance to trimethoprim, *oqxAB, qnrB, qnrS* among those producing resistance to quinolones, or *vanB,vanA* for glycopeptides-resistance). ResCap also detected more genes conferring resistance to heavy metals (e.g. cadmium, copper, silver or mercury), and relaxases, which are markers of plasmid families that carry antibiotic resistance genes (MOBC, MOBF, MOBP1, MOBP2) (**Figures S5-S7**).

### Comparative analysis of resistomes from different samples

A statistical analysis of “gene abundance”, analogous to that used for comparing the abundance of mRNA among samples in differential expression analysis^27^, was performed to quantify the improvement of ResCap over MSS in samples from different hosts. The need for such comparisons is based on the known differences in microbiotas of different hosts.

“Gene abundance” data without normalization were processed as “count data” and used as “input data” for differential analysis of the genes (detection of the genes only present in either human or swine samples) (**
Figure 5
**). Using MSS, the resistome of the total sample analyzed comprises 88 MGCs differentially detected (60 MGCs from humans and ^28^ MGCs from swine) with a p-value lower than 0.001. Conversely, ResCap detected 262 MGCs (186 from humans and 76 from swine) (**Figure 5, panel a**). Out of these 262 MGCs, 185 were differentially detected by ResCap and not by MSS, 77 were differentially detected by both approaches and 11 MGCs were only differentially detected by MSS (**Figure 5, panel C**). This result means that ResCap detected roughly three times more the MGCs on each resistome than MSS. The 11 MGCs that were only detected by MSS belong to common (“present in both human and swine samples”) MGCs by ResCap, suggesting that these differentially detected MGCs might in fact represent false positives. Meanwhile, the number of common MGCs detected in human and swine sets was 437 with MSS and 569 with ResCap, of which 269 MGCs were disclosed by both approaches, 300 MGCs being specific for ResCap and 168 for MSS. The 168 MGCs detected as common between human and swine metagenomes with MSS but not with ResCap were identified to be differentially present by ResCap as false negatives. This can be explained because the count of reads by MSS is lower than that of ResCap which makes the statistical analysis confidence values by ResCap better for a given MGC.

**Figure 5.**
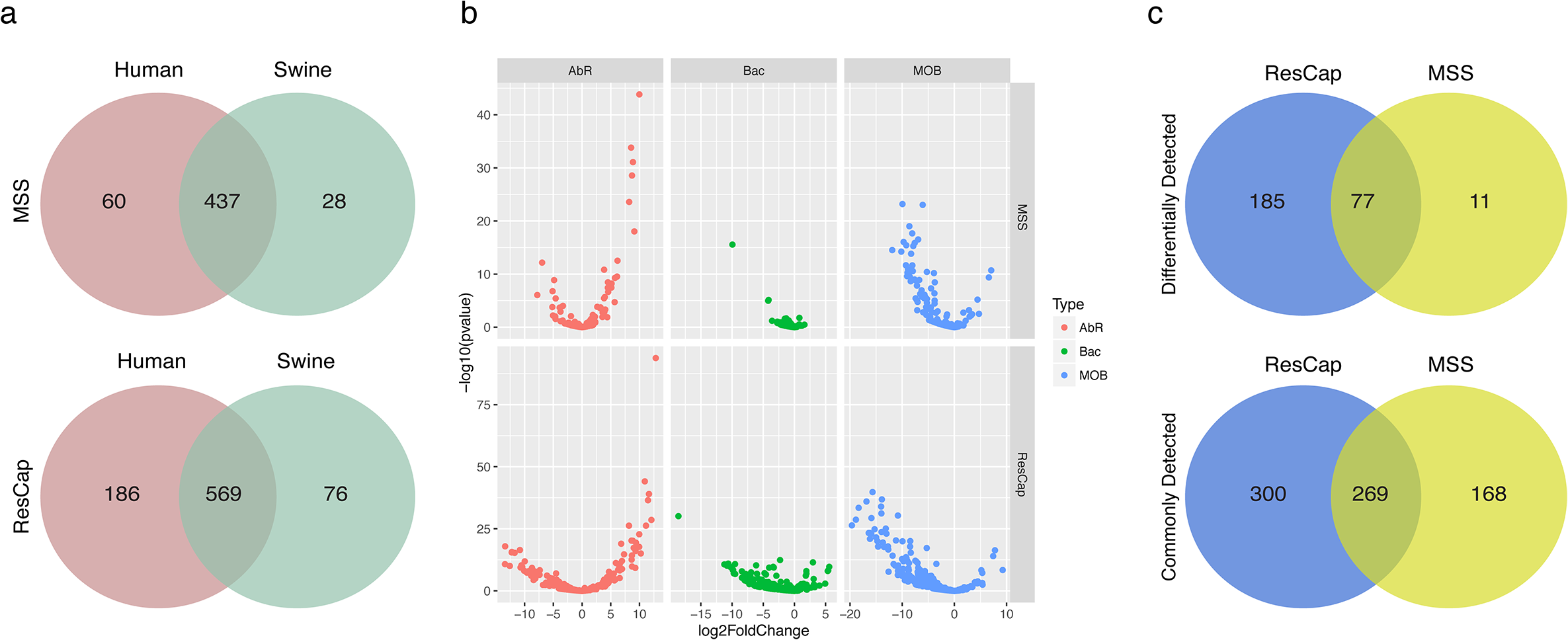
Differential study plots. Panel a) summarize the number of statistically significant MGCs of humans, swines and the genes in common between them using both approaches: MSS (up) and ResCap (button). Panel b) show the distribution of abundance variation between swine and human AbR resistomes (left), Metal and Biocide resistome (middle) and mobilome (right) in the form of volcano plots (fold change *vs* p-value) using the different approaches MSS (up) and ResCap (button) Left and right branches in the volcano refers to higher abundance in humans and swine respectively. Panel c) shown the Venn diagrams between approaches of differentially detected MGCs (up) and commonly (in both sets) detected MGCs (bottom)

**Figure 6.**
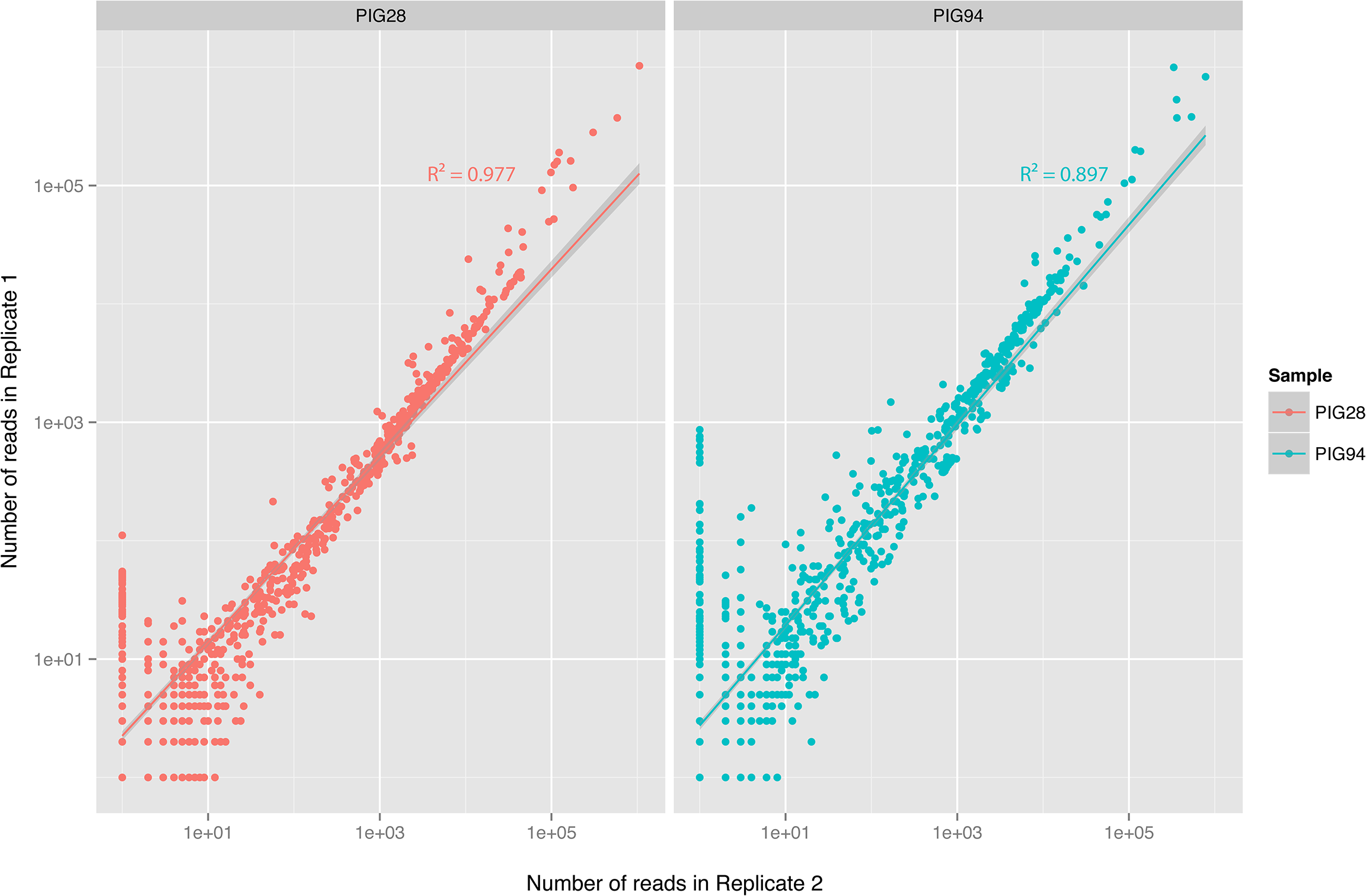
Reproducibility of ResCap. Reads from replicates are represented in dot plot to illustrate the linearity of the results from ResCap sequencing. Dots represent the genes detected in any of the replicates. Pearson’s product-moment correlation was used to estimate the correlation between technical replicates.

### Reference-free evaluation

ResCap includes approximately 78,600 genes that are homologous to “known” resistance genes, with different degrees of sequence identity with defined resistance genes, which might be involved in antibiotic resistance.

Assembly statistics and coverage show that the information obtained with ResCap only covers a small portion of the metagenome, as intended by design (**Figure 7**). As expected, ResCap increases, with respect to MSS, the number of sequenced genes that are homologous (evolutionary close) to the canonical genes included in Arg-ANNOT, BACMet and ConjDB databases. To perform a comparative analysis, the genes were catalogued as “ResCap”, “UniProt” or “Novel”. The “ResCap” gene set includes genes within the ResCap database of canonical genes. The “UniProt” gene set comprehends those that are already described in UniProtKB database and result in a positive blast against ResCap database. The “Novel” gene set corresponds to those genes not included in UniProtKB but resulting in a positive blast against ResCap canonical database. Only Blast hits with e-values lower than 10-100 were considered as positive and included in the analysis.

**Figure 7.**
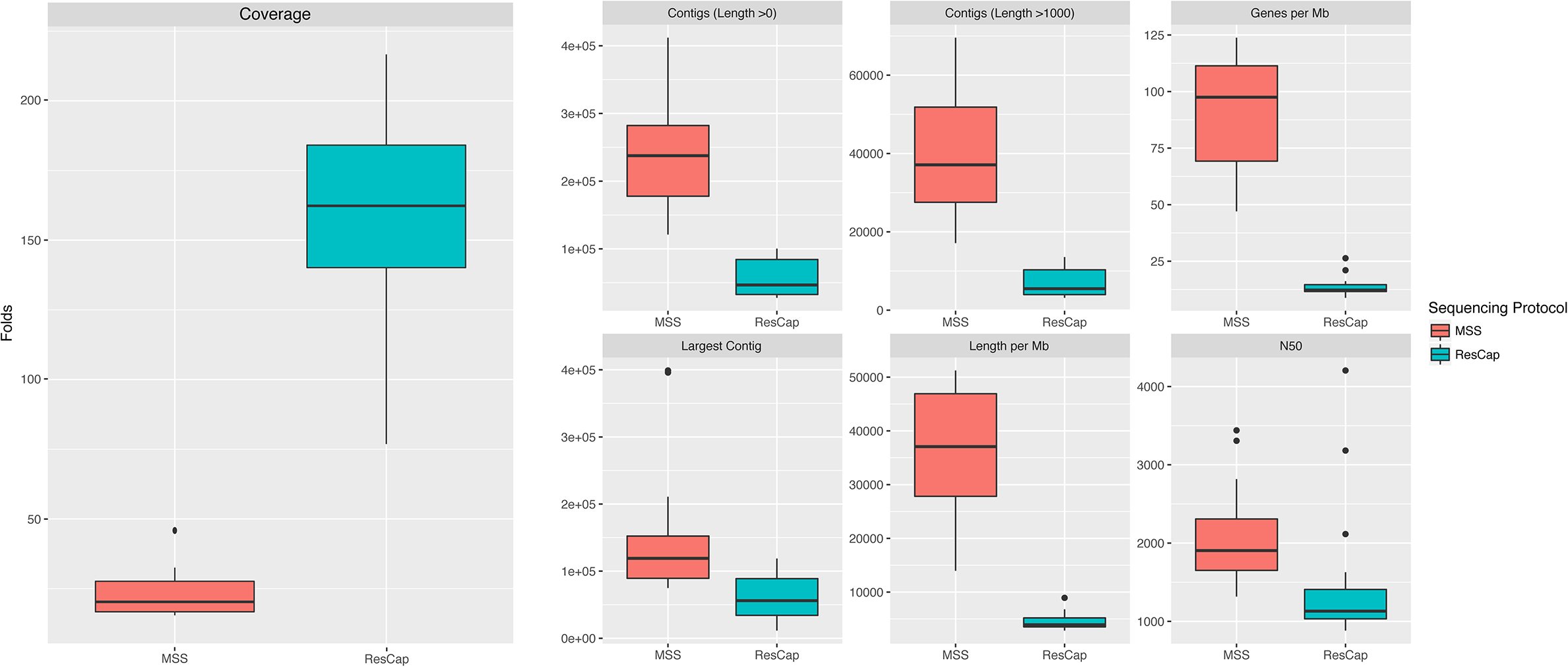
Assembly statistics. Statistic summary of the main assembly variables. *Total Length* and *Numberof genes* were normalized by the total amount of megabases sequenced by each sample. Coverage data was calculated as the total sequenced bases divided by the total length (without normalizing). Assembly statistics was calculated by Quast software.

The annotation of the genes shows that ResCap also improves the recovery of genes homology with genes coding for resistance, (UniProtKB 752±237 genes with ResCap *vs* 237±107 for humans and 441±71 genes vs 82 ±46 for swine with MSS; Novel genes, 79±38 genes with ResCap *vs* 20±7 107 for humans and 105±26 genes vs 9±4 for swine with MSS) as presented in **
Figure 8.** Although the actual role of these genes in antibiotic resistance will require functional validation that is beyond the scope of the current study, its identification as *bona fide* resistance genes as well as the analysis of their abundance upon antibiotic challenge might have a deep impact in further studies on the evolution of antibiotic resistance. **Figure S8** shows the better resolution of ResCap expressed by number of blast hits per gene per megabase.

**Figure 8.**
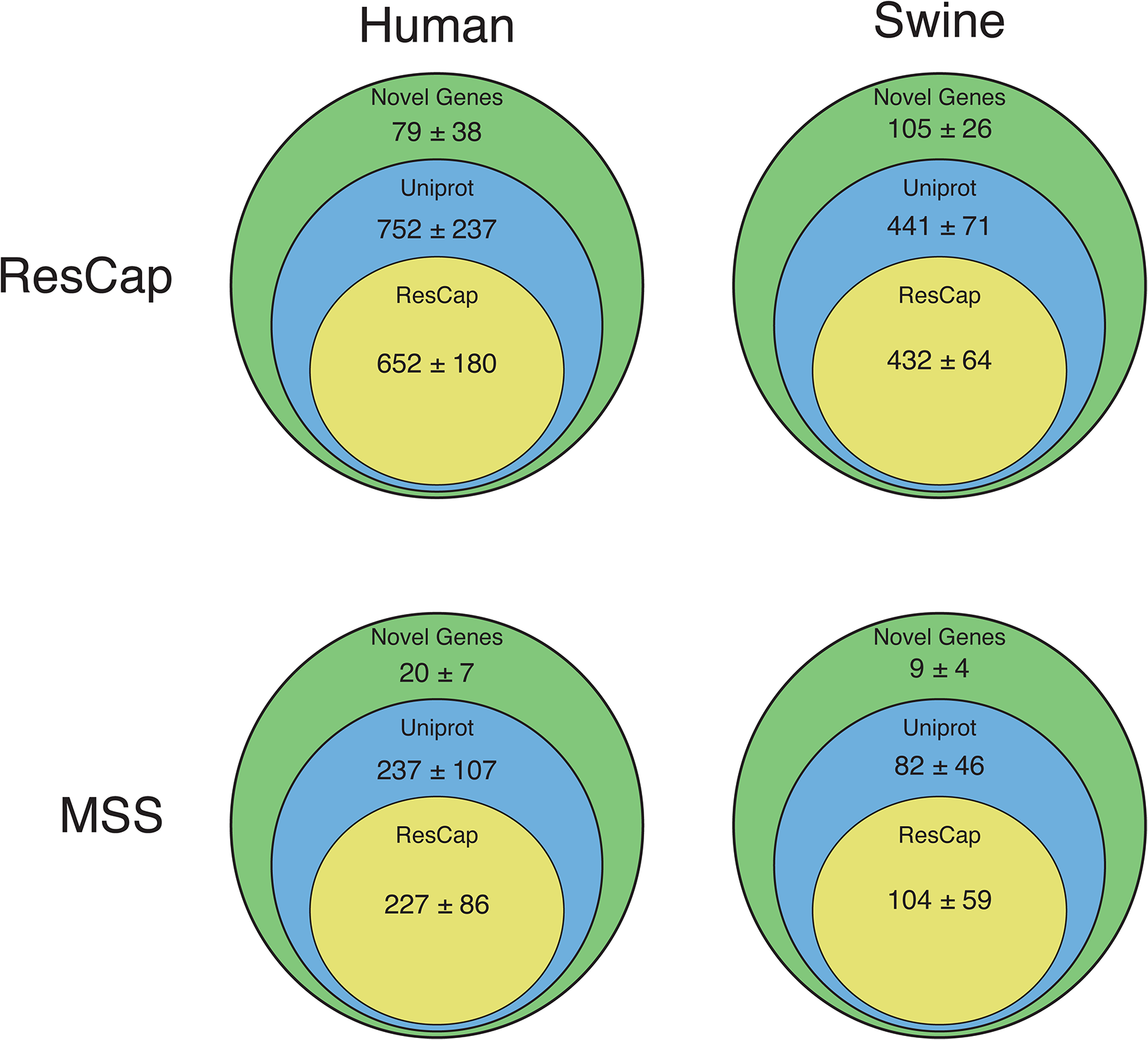
Functional annotation distribution. Assembled genes are classified as ResCap, UniProt or Novel (see Material & Methods). All assessed genes have a maximum e-value of 10^-100^ with some of the genes included in ResCap database. Figure show the comparative between human and swine samples and between MSS and ResCap approaches.

## DISCUSSION

This work reports the development of a novel resistance gene capture platform ResCap and on its comparative evaluation with MSS in resistance gene identification in a collection of human and swine faecal samples. Our study shows that ResCap is ideally suited for high-resolution analysis of resistomes and also offers the possibility to detect genes homologous to “known” resistance genes to further analyse the evolution of antibiotic resistance.

ResCap also provides several technical advantages to study resistomes in comparison with current metagenomic methods. First, the enrichment of ResCap resides in its targeted metagenomics approach, which significantly increases the recovery of sequences of resistance genes. Thus, ResCap reduces the sequencing depth needed to comprehensively detect the targeted genes and, consequently, contributes to lower sequencing costs. More importantly, it can significantly lower the limit of detection of resistance genes in complex microbiomes. Our results indicate that the resistome represents barely 0.2% of the gut metagenome. As a consequence, MSS needs at least 3.75×109 reads per sample to reach a similar coverage to that obtained by using ResCap (average of 1.9 × 107 paired reads that represents a relative enrichment of 279x). Second, the tiling of capture probes greatly facilitates the higher level of “gene horizontal alignment” coverage of ResCap relative to MSS (**Figure 2, Figure S2**), thus increasing specificity (**Figure S9**). Third, ResCap ability to detect previously unrecognized DNA fragments with homology to canonical resistance genes will facilitate the discovery of novel genes potentially involved in antibiotic resistance. In case they are antibiotic-selectable, the novel genes will be enriched in the presence of antibiotics. In addition, ResCap will be of interest in Public Health, because it allows a more accurate “ranking risk analysis”18 of the genes within the resistomes of different microbiotas. Finally, the substantial capacity of the platform (200Mb) makes ResCap extensible up to two fold of its current capacity, thus making possible its updating with new sequences published or added to resistance gene databases. ResCap updates will be publicly available through the GitHub repository and the Nimblegene webpage. Nonetheless, the threshold of detection of ResCap remains unknown due to the lack of a negative control that demonstrate the ability of ResCap to pick antibiotic resistance genes from quantified minority populations (e.g. mock genomic populations). Although appropriate, the complexity and variability of the metagenomic samples makes difficult to use a good negative control to this kind of studies.

The definition of parameters that accurately express antibiotic resistance “gene abundance” and antibiotic resistance “gene diversity” constitute a requirement to comparatively analyse the resistomes of different samples. Relative abundance parameters are widely used in computational analysis of MSS datasets, but require specialized statistics, as these compositional parameters are influenced by the variability in metagenomes of different samples. The application of the novel concept of MGCs (Mapping Gene Clusters, groups of alleles detected by the same set of reads) provides a set of normalized variables that can be measured in abundance and diversity among samples. Furthermore, the MGC-based system permits to evaluate the diversity within and between different functional groups (in our case, families of antibiotics, groups of genes conferring resistance to heavy metals or biocides and plasmid relaxases). To date, only a very few quantitative metagenomic approaches to analyze resistomes are available but they do not achieve this level of accuracy^14,16^.

Because of its sensitivity, specificity, and the possibility to accurately compare results between samples, ResCap complies with the needs of public health epidemiology of antibiotic resistance that include: i) the detection of emerging antibiotic resistance risks in different microbial environments^28^; (http://www.efsa.europa.eu/en/press/news/140325) ii) the need for implementation of accurate risk assessment studies based on resistome analysis in healthy humans, hospitalized patients, animal husbandry, food industry, and the environment; iii) the quality control of sewage and water bodies decontamination of antibiotic resistant genes iv) the update and refining of the list of resistance genes to be considered in monitoring the adverse effects of drugs in microbiomes, including pharmacomicrobiomic applications in clinical trials; v) the close monitoring of the efficacy of microbiome reconstitution/re-biosis, whether through targeted probiotic-live culture administration or fecal microbiota transplantation, to alleviate the adverse impact of antibiotic administration, and vi) to analyse the effect of eco-evo drugs and strategies to combat antibiotic resistance^29^.

In summary, ResCap provides an opportunity to meet the challenge of analyzing samples with complex and heterogeneous mix of genes in low and high concentration DNA samples. Thus, ResCap-like approaches might also be used to other complex microbial systems and their minority bacterial populations (e.g. virulence determinants, key-ecological traits involved in biosynthesis or biodegradation, or relevant genes of biotechnological interest).

## METHODS

### ResCap design

The ResCap capture library was based on a homemade core reference database (it will be available as per request) that comprises both well–known and hypothetical genes encoding resistance to antimicrobials (antibiotics, heavy metals, biocides) and genes coding for plasmid family markers (relaxases). The core reference database was built by downloading sequences associated with non-redundant antimicrobial genes available in curated databases Arg-ANNOT ^30^, CARD^31^, RED-DB, http://www.fibim.unisi.it/REDDB/Default.asp ResFinder^32^ and Bacmet ^33^.

The putative antibiotic resistance genes dataset was constructed as follows. All antibiotic resistance databases were combined in a non-redundant set. Proteins were clustered in protein families by homology, using CD-HIT with parameters of 80% identity and 80% coverage. First, each protein family was aligned by MUSCLE v. 3.7 ^34^ with default parameters. Then, a Hidden Markov Model (HMM) was built for each family with *hmmbuild* function of the HMMER3^35^ using default parameters. Hmmer search function (hmmsearch) was used against UniProtDB for each HMM profile to search homologous proteins for each family of proteins that confer antibiotic resistance. Manual curation of datasets was performed to remove false positives. Final protein data set was translated to DNA sequence using ENA accession numbers associated with each UniProtDB entry.

As a result, the final ResCap targeted sequence panel consists of 78,600 non-redundant genes (81,117 redundant genes) that would search a target space of 88.13Mb, not reaching yet the 200Mb target capacity offered by the custom SeqCap EZ library format (NimbleGen). Probes targeting the antibiotic resistome include 47,806 putative antibiotic resistance genes and 7,963 functionally characterized, canonical, antibiotic resistance genes. Probes targeting the metal and biocide resistome include 30,794 putative resistant genes and 704 canonical resistance genes. The platform also includes probes for 2,517 relaxases of the Conj database.

The consolidated list of target sequences was submitted to Roche NimbleGen for capture library design and synthesis and further implemented under the custom NimbleGen SeqCap EZ Developer Library format. Redistribution of probes for better capture uniformity, redundancy, and comprehensive target base coverage relied on NimbleGen, and was based on patented algorithms. ResCap design covers the 98.3% of the 88.13Mb and 99.6% of the genes have more than 50% of their sequence covered. (**Figure S9**).

### The ResCap workflow

The Rescap workflow consists of: i) whole-metagenome shotgun library construction, ii) hybridization, and iii) capture. All steps were performed according to NimbleGen standard protocols for Illumina platforms. To evaluate ResCvariable that would make it possibleap efficiency, samples were sequenced before and after capture.

i. *<u>Whole-metagenome shotgun library construction</u>*. Total nucleic acid was extracted following the standardized Metahit protocol^36^ (http://www.metahit.eu/ and using the FastPrep instrument (MP Biomedicals, USA). Libraries were prepared following the instructions of “Kapa Library Preparation Kit for Illumina platforms” (Kapabiosystems, KR0935-v1.13). Briefly, 1.0 µg input DNA (measure by Picogreen) was fragmented to 500-600 bp insert size by sonication with Bioruptor (FastPrep®-24). After End repair, A-tailing and Adapter ligation, we follow Dual-SPRI size selection adding 0.5 vol in first cut and 0.2 vol in the second cut to get 650-750pb libraries. Library amplification was carried out using LM-PCR of 7 cycles, as indicated in the SeqCap EZ Library SR User´s Guide v4.2. At this level, samples were labelled with specific barcodes for further sample identification. A first aliquot of the resulting amplified libraries were quality checked in a Bioanalyzer 2100 (Agilent) and pooled in equimolecular amounts for sequencing on Illumina HiSeq 2000 instrument, generating 100-150-bp paired-end reads (“pre-capture” samples).
ii. *<u>Hybridization and capture</u>*. The second part of each DNA library was subjected to targeted sequence capture with the custom ResCap probes prior to sequencing (“post-capture” samples). Both experiments were made in separate sequencing runs. Targeted sequence capture was carried out according to the manufacturer’s specifications. The captured DNA was checked for quality and integrity in a Bioanalyzer and titrated by quantitative PCR using the “Kapa-SYBR FAST qPCR kit forLightCycler480” and a reference standard for quantification. The captured libraries were denatured prior to be loaded on a flow-cell at a density of 2,2pM, where clusters were formed and sequenced using a HiSeq 2000 in a 2x100 pair-end mode for swine samples and NextSeq 500 in a 2x150 pair-end mode for human samples. Raw sequences were processed using FastX Toolkit (http://hannonlab.cshl.edu/fastx_toolkit/).

### Bioinformatic analysis

#### Reference-based workflow

Analysis of sequence data from metagenomes constitutes a challenge because of the inherent variability of the samples analysed, and the limitations of current bioinformatics’ methods to unequivocally identify specific alleles from short length reads (100 −150 bp). To overcome such limitations, we developed a novel approach to define variables suitable for inferring “gene abundance” and “gene diversity” and, in our case, to perform quantitative analysis of antimicrobial resistance genes. Moreover, we suggest a workflow of variable normalization in relation to the information content of the targeted variable that would make it possible to compare different samples of different hosts. These tools were developed for ResCap but could be implemented for any other metagenomic sequence dataset. Shotgun metagenomic sequencing allowed assembling the sequences into contigs to infer the functionality of the sequenced metagenome. **Figure 9** shows the workflow that illustrates and defines the variables used.

**Figure 9.**
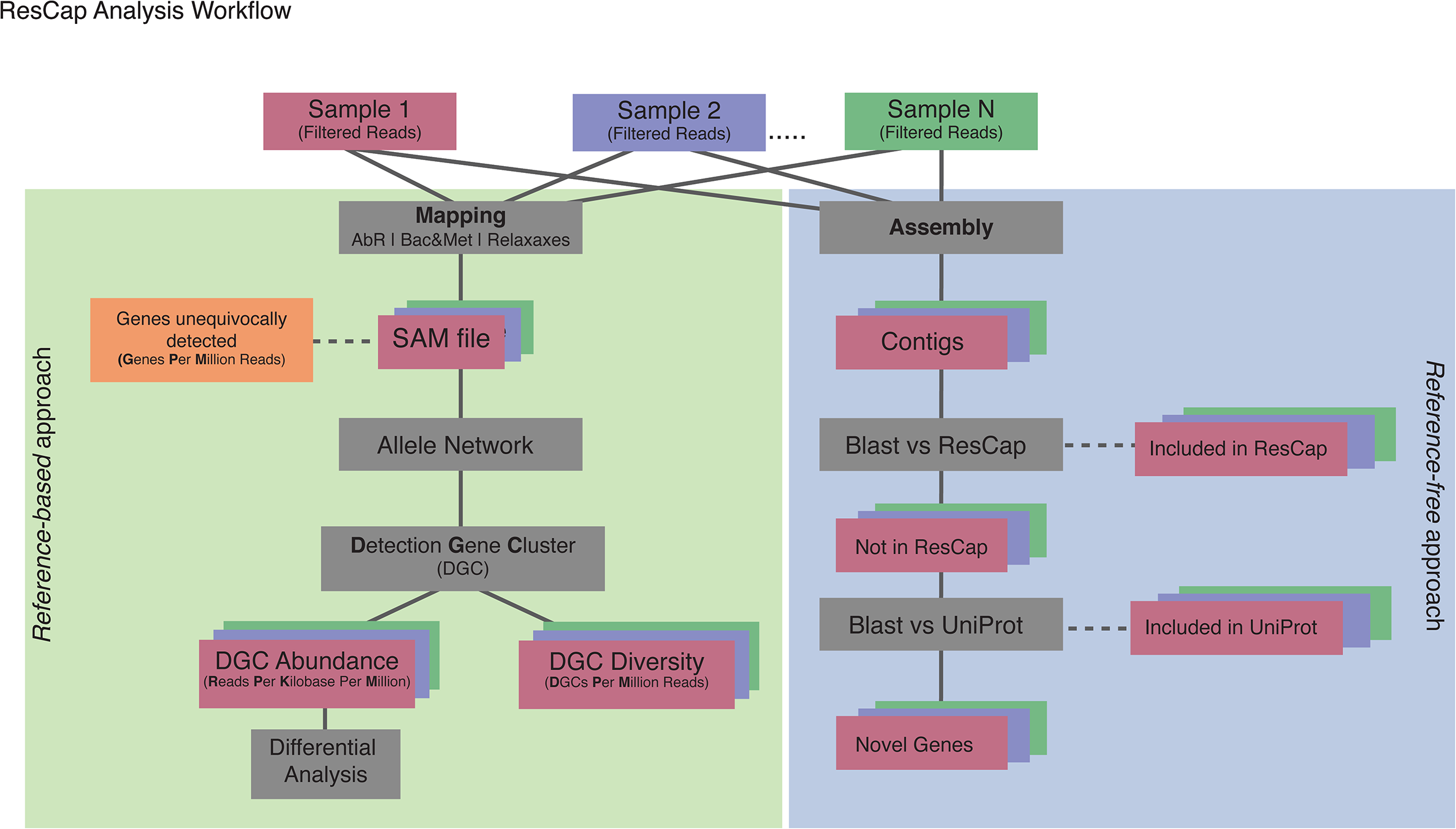
ResCap analysis workflow. Processed reads are mapped against reference database, SAM files are parsed to extract the reads unequivocally mapped and the ambiguously mapped to determine the Genes unequivocally detected and to perform the Allele Network. Allele Network is build using all SAM files of the study. The MGCs determines from Allele Network are used to perform the statistical analysis of Abundance and Diversity. Finally with the data of Abundance a differential analysis was performed.

##### Raw Data Processing

Reads were mapped against our database comprising ARG-ANNOT, BacMET and ConjDB databases independently, using Bowtie2 software^37^. Bowtie was set up to retrieve all end to end possible alignments and suppress both discordant alignments and mixed alignments. The output SAM file was parsed to get the fields of *Query template NAME, Reference sequence NAME,1-based leftmost mapping Position, MAPping Quality, Position of the mate/next read.* Reads with unavailable information (field *Query Template NAME* equal to ‘*’) were suppressed. Subsequently, a homemade perl script (available per request) was used to count matched reads per gene. Using the SAM parsed file and the length of the reference genes, the perl script generated a table with the following fields i) the number of reads per gene mapped (RPG, “gene depth coverage”), ii) the number of reads per kb of gene (RPK), iii) the number of the reads that were mapped unequivocally to a given gene and iv) the percentage of coverage of the gene sequence (“gene horizontal alignment coverage”) of each mapped gene. Table fields Unique, RPG and RPK were normalized by the total amount of reads in each sample, transforming such fields in “read per gene per million reads” and “reads per kb per million reads” (RPKM), respectively, the last one being a common unit of “gene abundance”^38^. Several ways to normalize abundance data have been applied to different studies (e.g., expression data in RNA-Seq experiment). The aim of our approach was to estimate the proportion of antimicrobial resistance genes among samples that putatively contained the same amount of DNA, so the normalization using the total amount of DNA (i.e reads) among samples fits better with the initial approach.

The redundancy of mapped reads may be represented as a network where the nodes are the genes (usually alleles of the same gene) and the edges are the reads that map in the different nodes. Because one read can map in different alleles/genes, all the genes mapped by these reads are linked among them. The resulting network that comprises all the nodes and edges in a set of samples is named “allele Network*”* (**Figure S10**). In our context, the allele network must be unique for all samples of a given assay, so an allele network was built joining all the SAM parsed files of the study.

Each cluster of the Allele Network represents the set of genes that are detected by a set of reads. They are defined as a Mapping Gene Clusters (MGCs) and each one may include hundreds of genes or just one gene. A given MGC will be detected when at least one read maps against any of the genes within that MGC (diversity). To quantify the MGCs in each sample, the highest value shown by an allele (node) within a given MGC is taken as the occurrence of such MGC (abundance). The MGCs system builds a set of normalized variables that can be measured in abundance and diversity among samples and thus, allows comparing datasets of different sources, while maximizing the accuracy of the observable information.

A homemade perl script was used to build the allele network from the SAM parsed files, taking the mapped genes as nodes and searching the ambiguously mapped reads to create the edges. Perl script calculates the edges-weight as the number of reads that map the linked nodes at the same time. Allele Network was loaded in R environment^39^ using the *igraph* package^40^. MGCs were defined using *mcl*, from MCL R package^41^, with default parameters except allow loops and cluster with only one member on the allele network.

##### Data Analysis

The resistome of a given experiment was analysed in terms of gene abundance and diversity according to the methodology described above. The *abundance* and the *diversity* of genes in a particular resistome are the (dependent) variables that define this resistome and are measured as the number of RPKM per MGC and the number of MGCs, respectively.

The number of MGCs was normalized by the total number of sequencing reads per each sample expressed in millions of reads (MPM), this value being considered as a unit of **diversity**. MGCs of the antibiotic resistance gene database were divided according to antibiotic families ^30^. MGCs of the relaxase database were organized in known different relaxase families ^42^. The MGCs of biocide and heavy metal resistance gene database were classified according the susceptibility to specific compounds ^33^. Genes that belong to more than one functional category (e.g. some conferring resistance to different metals) contribute equally for any of them. Figure S9 shows the 839 MGCs determined in our sample (237 for AbR, 283 for Biocide and Metals and 319 for relaxases). Descriptive statistic was performed using *dplyr*^43^, *tidyr*^44^ and *ggplot2*^45^ packages of R^39^.

Differential analysis was performed using DESeq2 package^46^. Although DESeq2 was originally designed for differential expression analysis, it also works well with abundance data. Tables containing the original abundance data obtained by ResCap and MSS datasets were used separately as input for DESeq2 package to determine the MGCs differentially detected between swine and human hosts. Normalization and statistical analysis were performed with the default parameters of DESeq2. MGCs with p-value lower than 0.001 were classified as differentially detected, rest of the MGCs (p-value above 0.001) were classified as commonly detected.

#### Reference-free workflow

Assemblies were performed by MegaHit software with default parameters^47^. Prodigal^48^ was used for gene recognition and translation with the specific parameters for metagenomic sequences. Quality assemblies’ quantification was performed by Quast software^49^. Predicted genes were first annotated against the ResCap database by Best Blast Hit approach using blastn software^50^. In order to identify only genes belonging to ResCap database or their homologs, and minimize the false positive ratio, Blast hits were filtered by e-value of 10 and 80% of coverage. Genes with identities higher than 95% and coverage higher than 80% were considered as belonging to ResCap. The remaining genes were translated to proteins. These proteins were classified as non-ResCap and were compared against UniProt by blastp. Again, hits with higher identity than 95%, coverage higher than 80% and e-value lower than 10^−100^ were considered as UniProt known proteins. The set of proteins that did not accomplish this threshold were considered as novel proteins.

### Samples analyzed

ResCap was validated by analysing fecal samples from 9 human and 8 swine individuals, all collected as part of FP7 European Research Consortium EvoTAR (www.evotar.eu). Swine samples were collected in Spanish farms linked to large companies which supply broilers and swine processed meat in the EU. Antibiotics as growth promoters or with preventive purposes are not used in these farms. Human samples were collected in the Hôpital Bichat, Paris, France, under the protocol approved by its local ethics committee. DNA preparation was accomplished for animal and human samples using standardized protocols (MetaHIT Protocol). Robustness of the platform was tested by comparative analysis of two technical replicates of two swine samples.

**Table 2.**
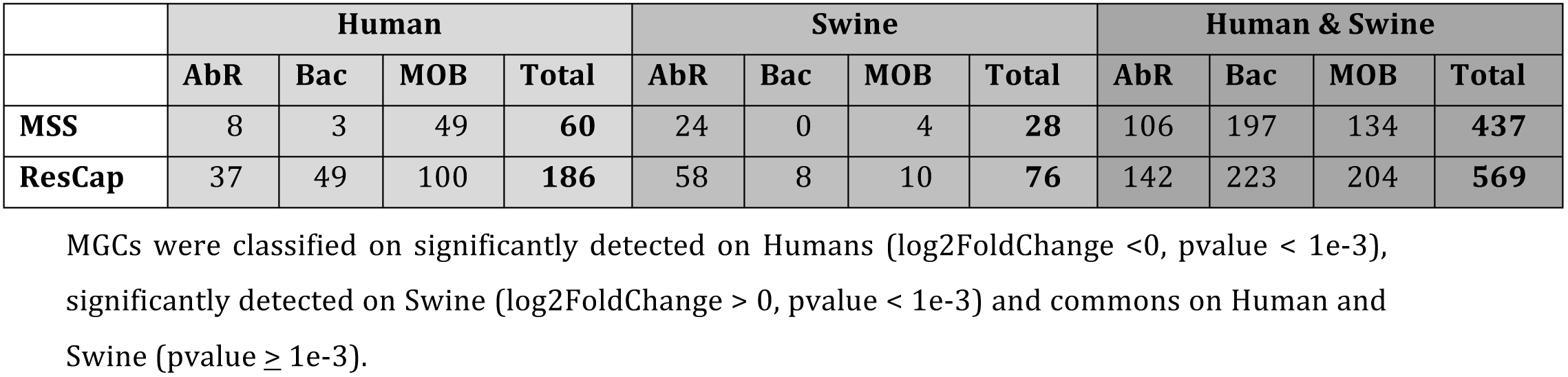
Summary of metagenomic comparative analysis

|  | Human |  |  |  | Swine |  |  |  | Human & Swine |  |  |  |
| --- | --- | --- | --- | --- | --- | --- | --- | --- | --- | --- | --- | --- |
|  | AbR | Bac | MOB | Total | AbR | Bac | MOB | Total | AbR | Bac | MOB | Total |
| **MSS** | 8 | 3 | 49 | **60** | 24 | 0 | 4 | **28** | 106 | 197 | 134 | **437** |
| **ResCap** | 37 | 49 | 100 | **186** | 58 | 8 | 10 | **76** | 142 | 223 | 204 | **569** |

MGCs were classified on significantly detected on Humans (log2FoldChange <0, pvalue < 1e-3), significantly detected on Swine (log2FoldChange > 0, pvalue < 1e-3) and commons on Human and Swine (pvalue ≥ 1e-3).

## ACKNOWLEDGEMENTS

This work was supported by the European Commission, Seven Framework Program (EVOTARFP7-HEALTH-282004 for VFL, FB, JLM, AA, DE, ER, RJLW, WvS, FdlC and TMC), the Join Programming Initiative in Water (JPI Water StARE JPIW2013-089-C02-01 to JLM), the Ministry of Economy and Competitiveness of Spain (BIO2014-54507-R to JLM, and PLASWIRES-612146/FP7-ICT-2013-10 and BFU2014-55534-C2-1-P for FdlC). Authors also acknowledge the European Development Regional Fund “A way to achieve Europe” (ERDF) for cofounding the Plan Nacional de I+D+ I 2012-2015 (BIO2014-54507-R to JLM, PI12-01581 to TMC and BFU2014-55534-C2-1-P for FdlC) and CIBER (CIBER in Epidemiology and Public Health, CIBERESP; CB06/02/0053 to FB) and the Spanish Network for Research on Infectious Diseases (REIPI RD12/0015 to JLM), and the Regional Goverment of Madrid (PROMPT-S2010/BMD2414). Val F. Lanza was further funded by a Research Award Grant 2016 of the European Society for Clinical Microbiology and Infectious Diseases (ESCMID). Conflict of Interest: none declared

## COMPETING FINANTIAL INTEREST

The authors declare no competing financial interests

## LEGENDS TO SUPPLEMENTARY MATERIAL

**Figure 1 Supplementary.**
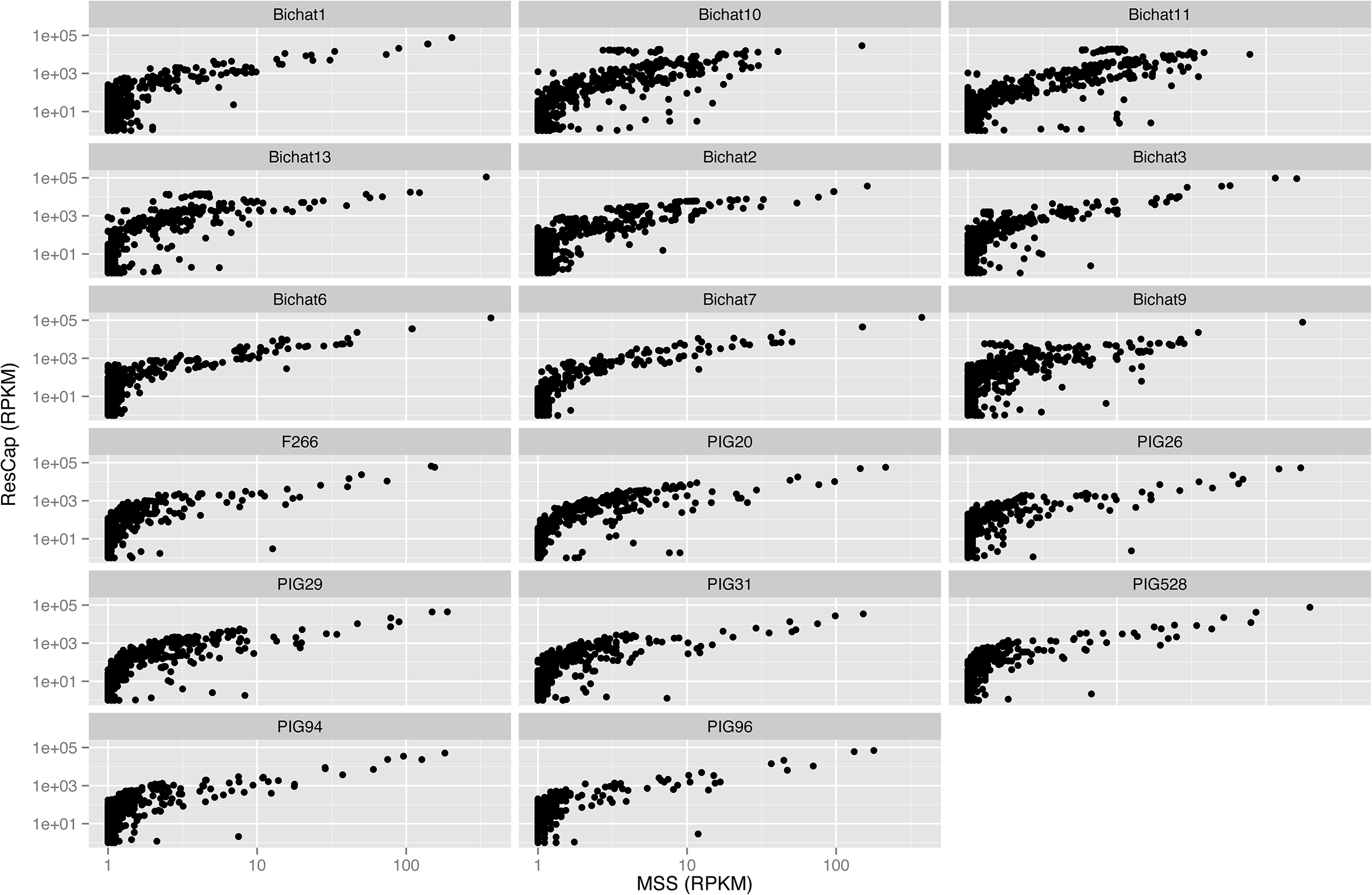
Gain function plot for each sample. Representation of the gain in reads per kilobase per million of reads of each detected gene between MSS (abscissa axis) and ResCap (ordinate axis). Genes only identified by ResCap are represented by the dot cluster in the initial values of abscissa axis. The pictures are represented in log-log scale to a better perception of the linearity of the gain function in genes detected by each protocol.

**Supplementary Figure 2.**
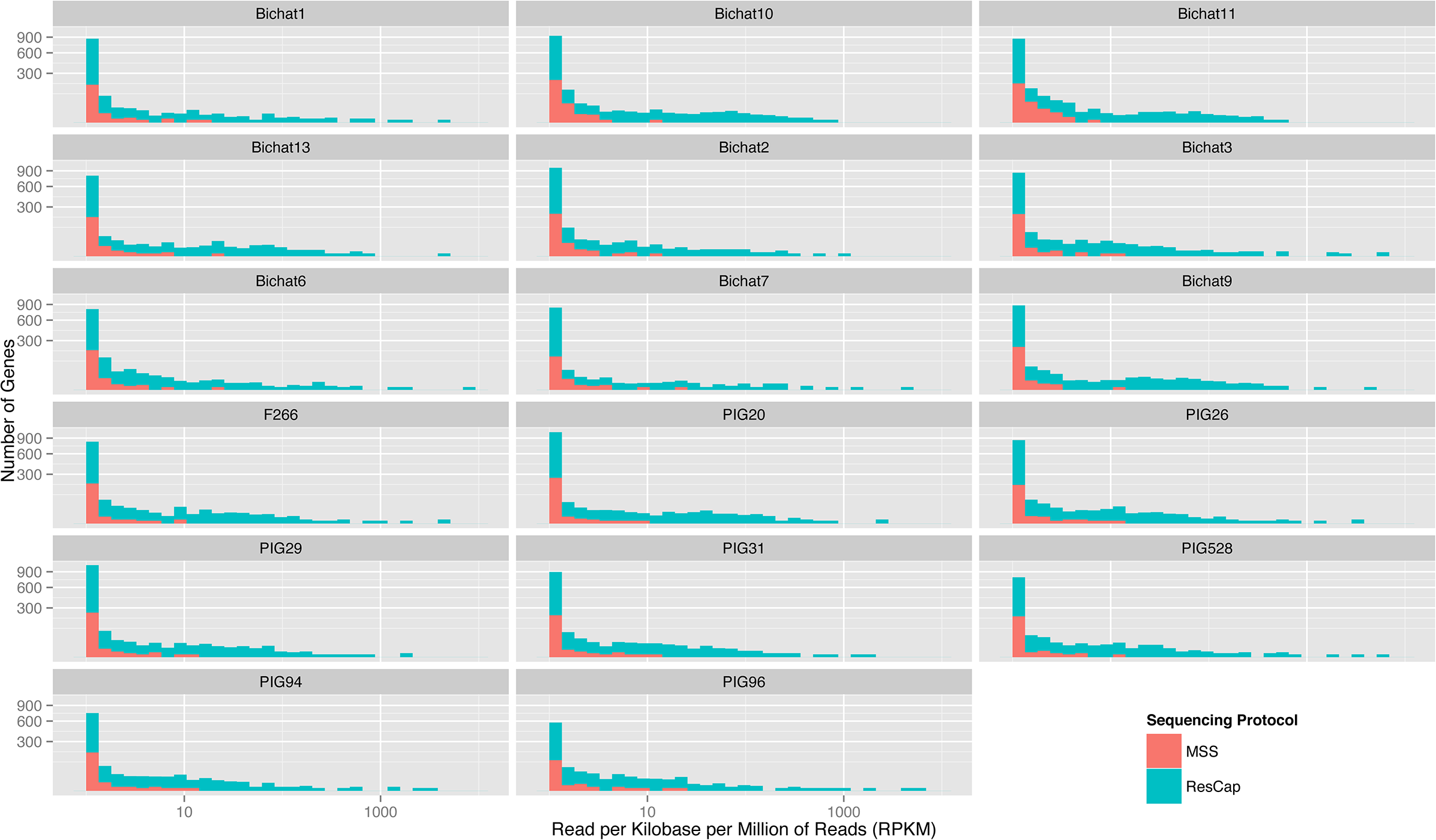
Distribution of Reads Abundance: Figure shown the histograms of reads abundance per each gene. Each frame represent a sample, superposing results from MSS protocol and ResCap protocol. A square scale was used for ordinate axis and a logarithmic scale for abscissa axis to optimize the representation of the data.

**Supplementary Figure 3.**
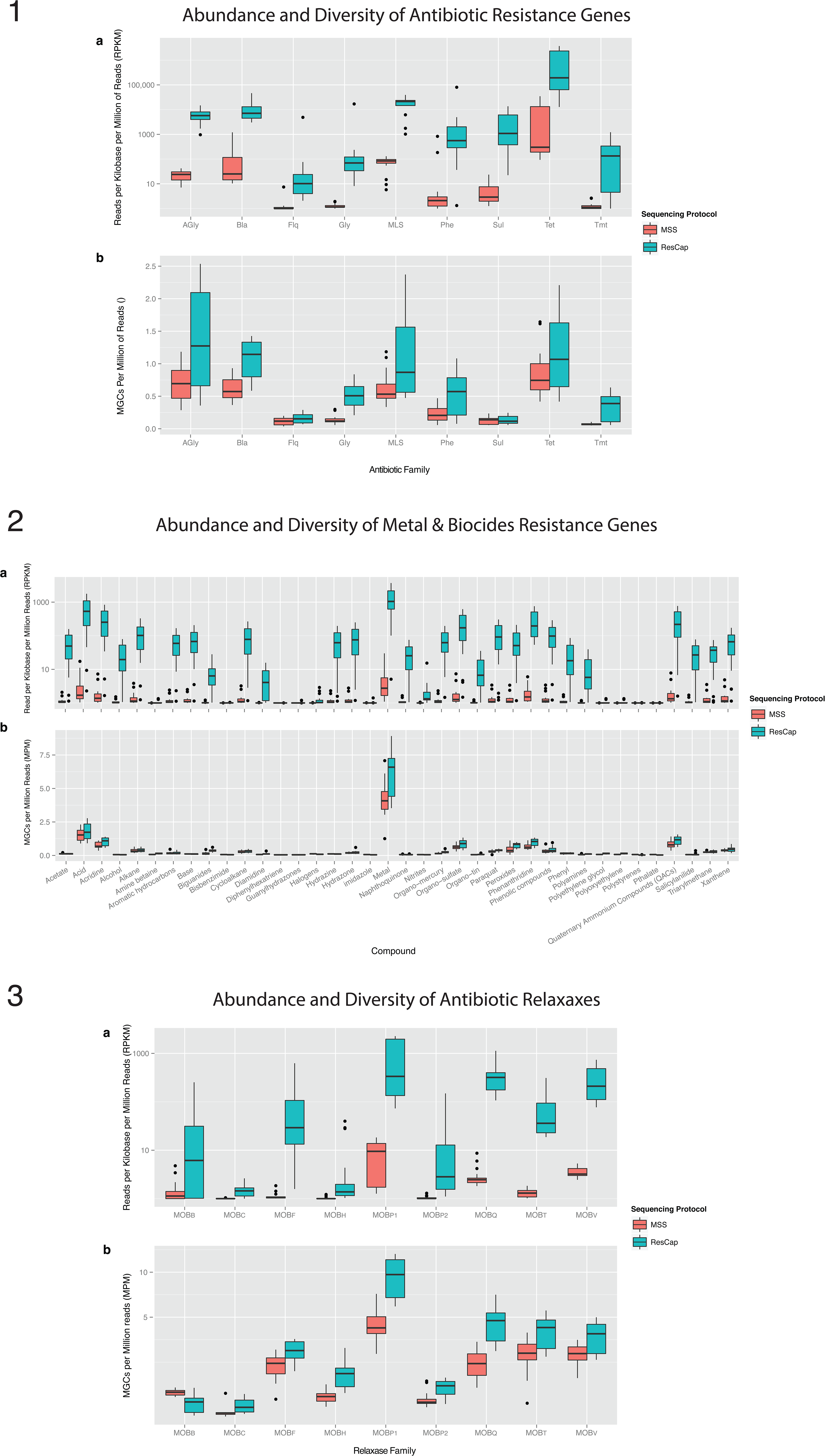
**(1) Diversity and Abundance of Antibiotic Resistance:** Comparison of ResCap and MSS protocol in Antibiotic Resistance data. Antibiotic resistance genes were divided among nine families by antibiotic family (AGly: Aminoglycosides, Bla: Beta-Lactamases, Flq: Fluoroquinolones, Gly: Glycopeptides, MLS: Macrolides, Phe: Phenicols, Sul: Sulphonamides, Tet: Tetracyclines and Tmt: Trimethoprim). Abundance (a) was measured as <u>R</u>ead <u>P</u>er <u>K</u>ilobase per <u>M</u>illion of reads that mapping against genes or allele-cluster genes of each antibiotic resistance family. Diversity (b) was measured as a number of detected Genes Per Million reads of each antibiotic resistance family. **(2)Diversity and Abundance of Relaxases:** Comparison of ResCap and MSS protocol in Relaxases data. Relaxases were divided among nine protein families (MOB_B_, MOB_C_, MOB_F_, MOB_H_, MOB_P1_, MOB_P2_, MOB_Q_, MOB_T_ and MOB_V_). Abundance (a) was measured as <u>R</u>ead <u>P</u>er <u>K</u>ilobase per <u>M</u>illion of reads that mapping against genes or allele-cluster genes of each relaxase family. Diversity (b) was measured as a number of detected <u>G</u>enes <u>P</u>er <u>M</u>illion reads of each relaxasa family. **(3) Diversity and Abundance of Biocide & Metal resistance:** Comparison of ResCap and MSS protocol in Biocide & Metal resistance data. Biocide & Metal resistance genes were divided by compound susceptibility. Abundance (a) was measured as <u>R</u>ead <u>P</u>er <u>K</u>ilobase per <u>M</u>illion of reads that mapping against genes or allele-cluster genes of each compound family. Diversity (b) was measured as a number of detected <u>G</u>enes <u>P</u>er <u>M</u>illion reads of each compound family.

**Supplementary Figure 4.**
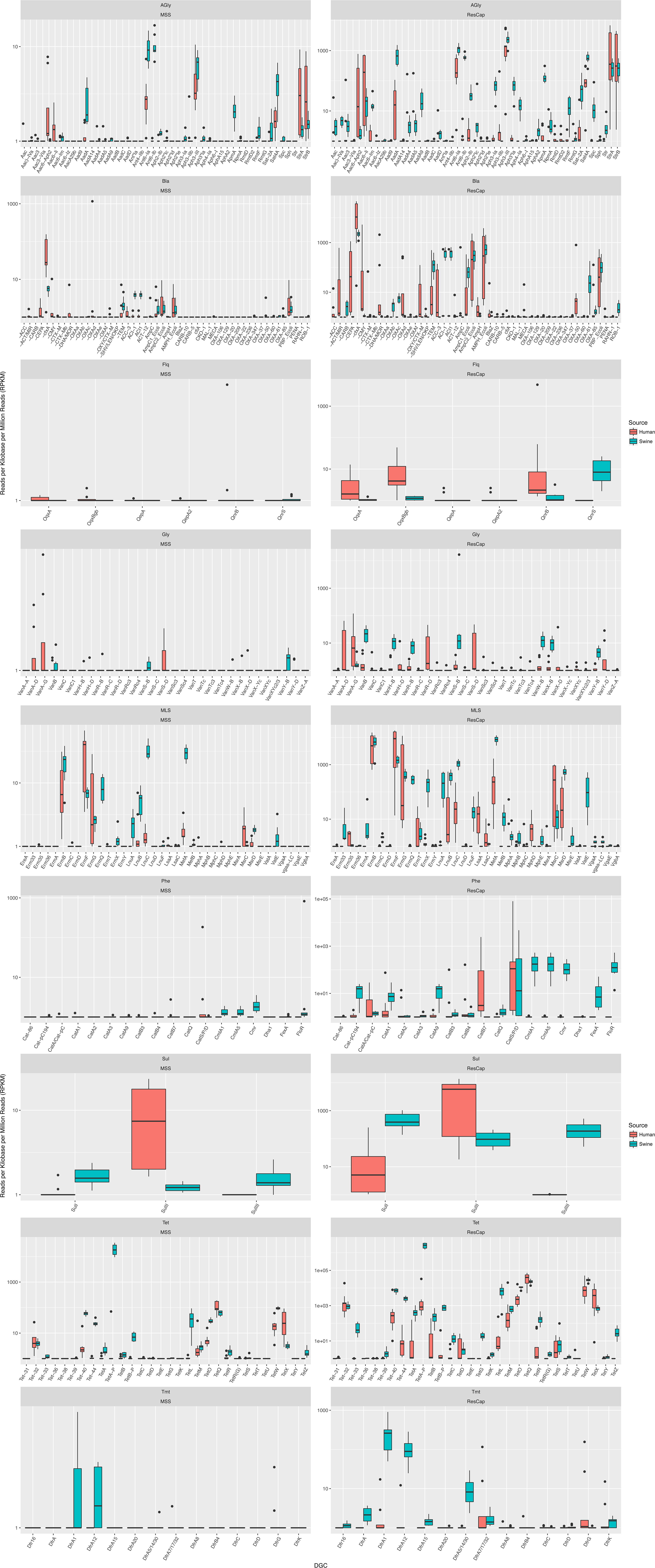
MGCs abundance comparative of antibiotic resistance between swine and human samples. MGCs corresponding to antibiotic resistance dataset were classified by antibiotic families (Agly: Aminoglycosides, Bla: Betalactams, Flq: Fluoroquinolones, Gly: Glycopeptides, MLS: Macrolides, Phe: Phenicols, Sul: Sulphonamides, Tet: Tetracyclines, Tmt: Trimethoprim). Abundance was measured as Read per Kilobase per Million of reads. Panel right shown the results of MSS and panel left shown the results of ResCap.

**Supplementary Figure 5.**
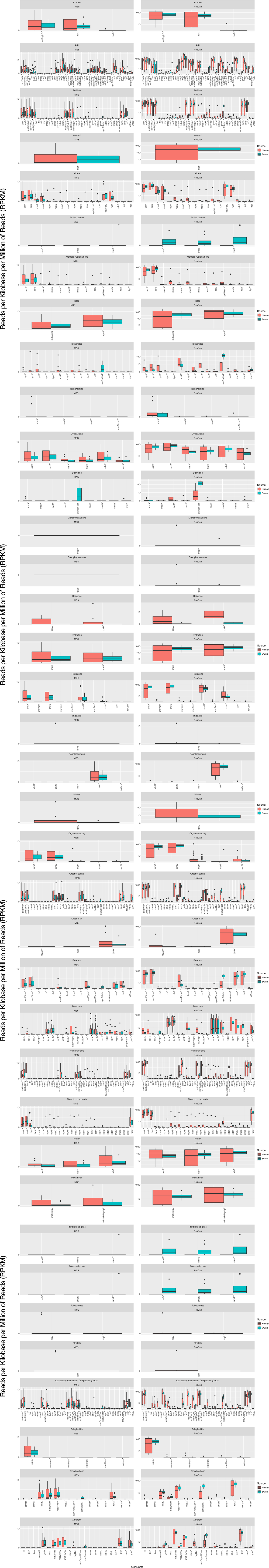
MGCs abundance comparative of biocide resistance between swine and human samples. Gene abundance was extracted from original count data after normalization. Some sets of genes make complex MGCs. In this representation, MGCs quantification was discarded in order to increase the biological information. Genes were classified by compound susceptibility. Due to biocide resistance genes spectrum of activity, genes are not constricted to one category but some genes show resistance to more than one compound. Genetic abundance is expressed as Reads per Kilobase per Million of Reads (RPKM). The panel right shows the results of MSS and the panel left shows the results of ResCap.

**Supplementary Figure 6.**
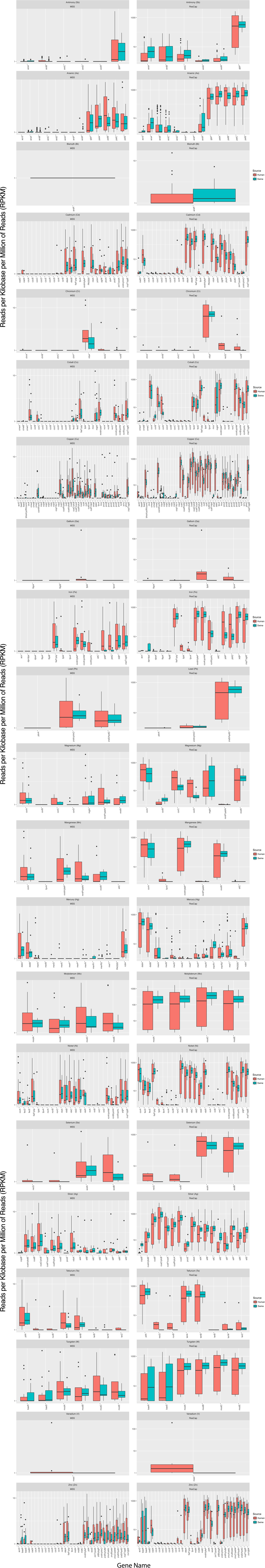
Gene abundance comparative of metal resistance between swine and human samples. Gene abundance was extracted from original count data after normalization. Some sets of genes make complex MGCs. In this representation, MGCs quantification was discarded in order to increase the biological information. Genes were classified by metal susceptibility. Due to metal resistance genes spectrum of activity, some genes are not constricted to one category but some genes show resistance to more than one metal. Genetic abundance is expressed as Reads per Kilobase per Million of Reads (RPKM). The panel right shows the results of MSS and the panel left shows the results of ResCap.

**Supplementary Figure 7.**
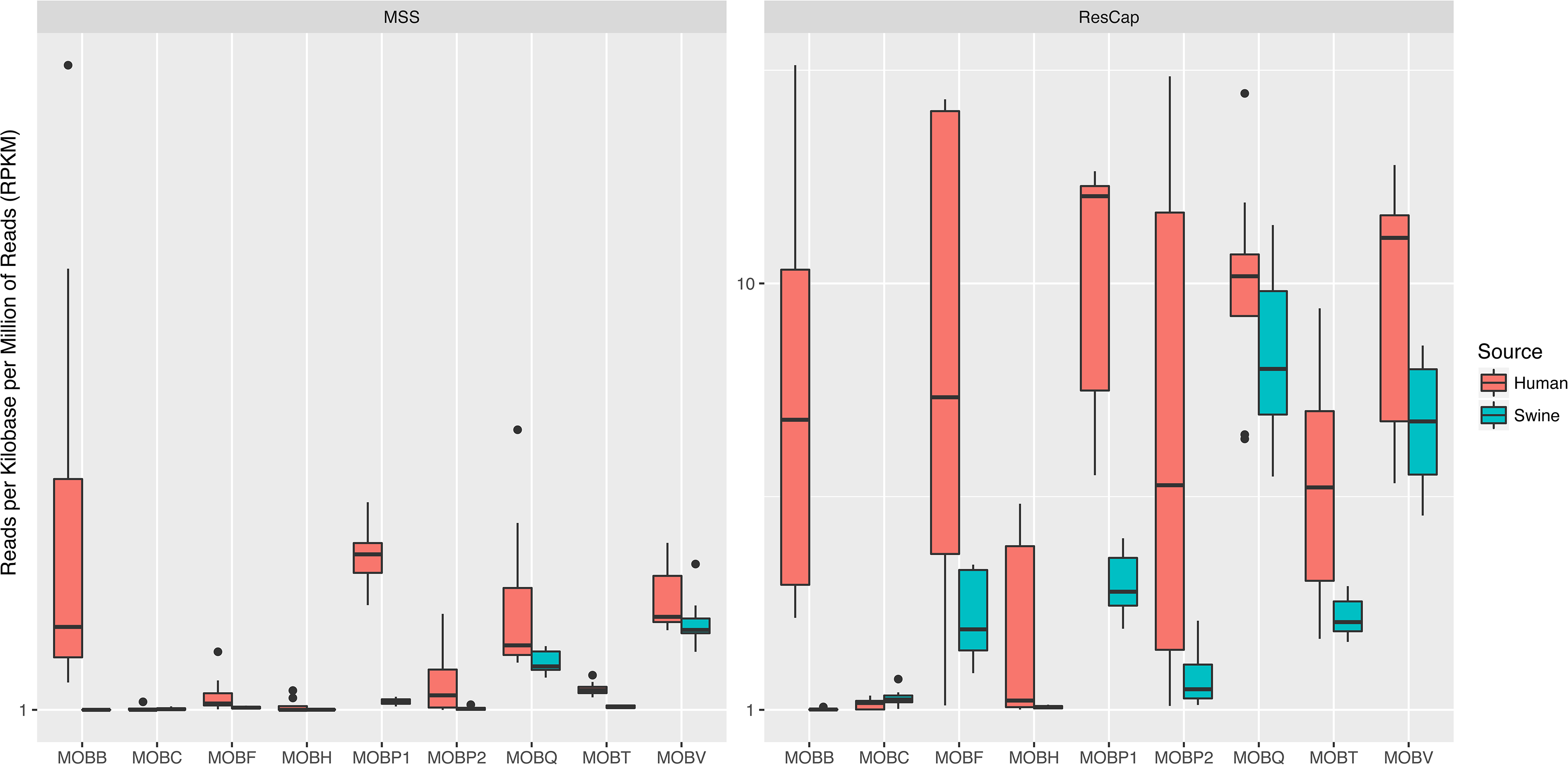
MGCs abundance comparative of Relaxases between swine and human samples. Relaxases were classified by MOB families. MGCs abundance was summarized in MOB families. Each MOB families are composed by several MGCs. Genetic abundance is expressed as Reads per Kilobase per Million of Reads (RPKM). The panel right shows the results of MSS and the panel left shows the results of ResCap.

**Supplementary Figure 8.**
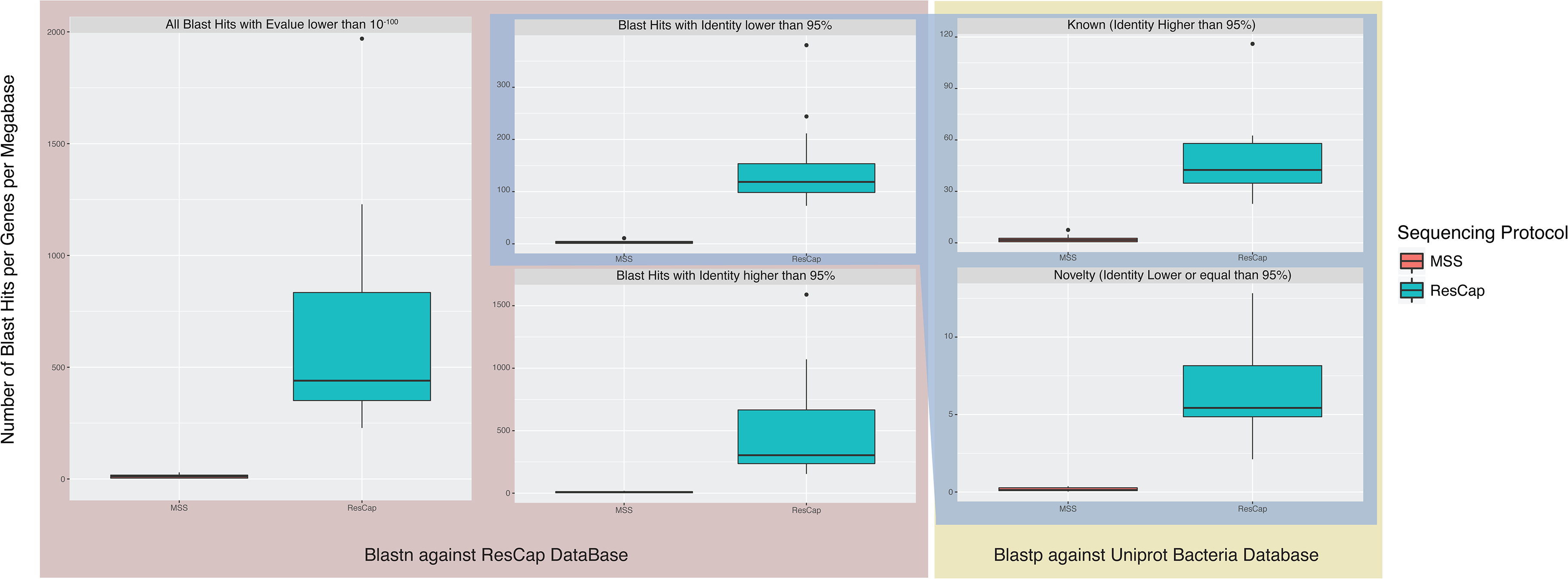
Blast annotations summary. Summary of the classification steps of assembled genes. The sequential annotation comprises a first blastn search for identify resistome homologous genes. Genes with evalue higher than 10^-100^ were discarded. Filtered genes were split into two groups, genes with identity higher than 95% and genes with identity lower than 95%. The second group were annotated against UniProtKB and were split again into two groups, genes with identity higher than 95% of identity and genes with identity lower of 95%. A number of blast hits were normalized by the number of assembling genes per sequenced megabases.

**Supplementary Figure 9.**
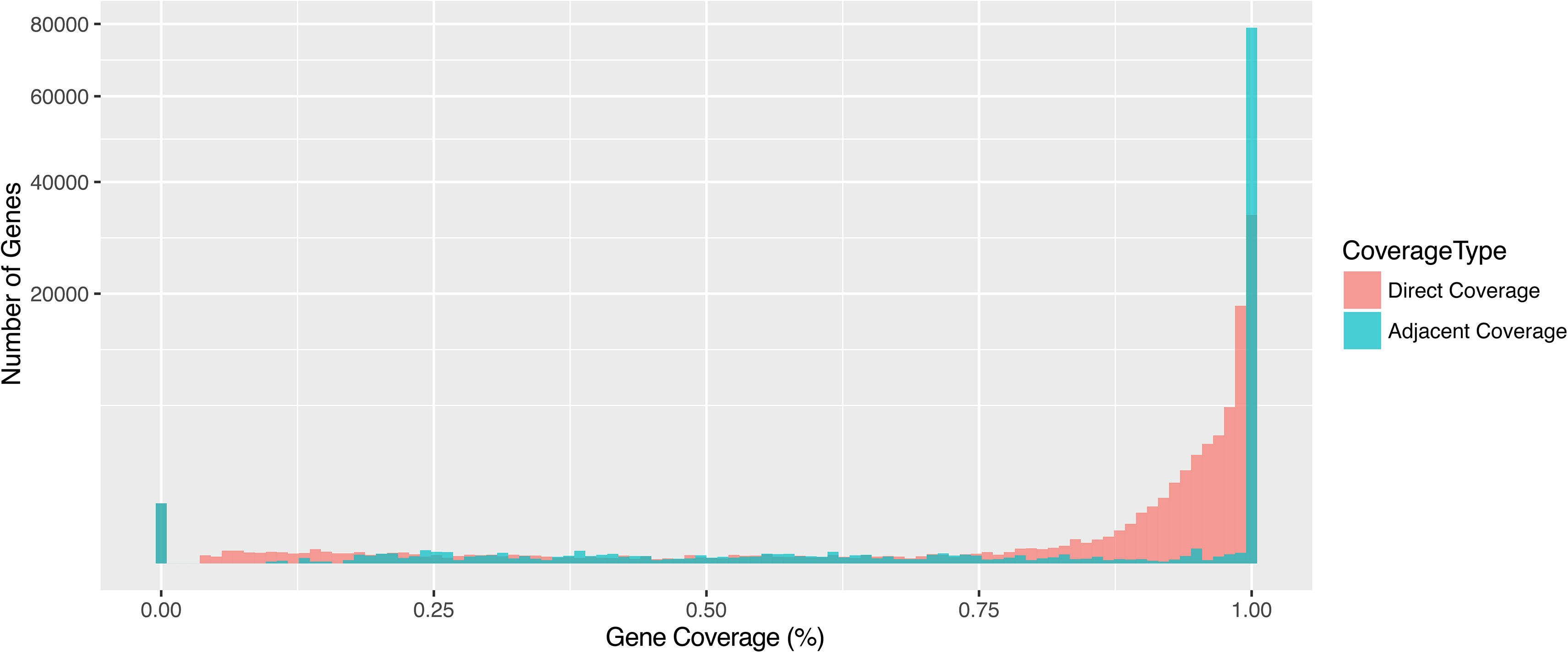
Histogram of gene coverage distribution by hybridizing probes. Two metrics was provided by NimbleGene, Direct Coverage (red bars) and Adjacent Coverage (cyan bars). 90% of the genes are covered at least 96.9% by direct coverage and 90% of the genes are covered at 100% of Adjacent Coverage.

**Supplementary Figure 10.**
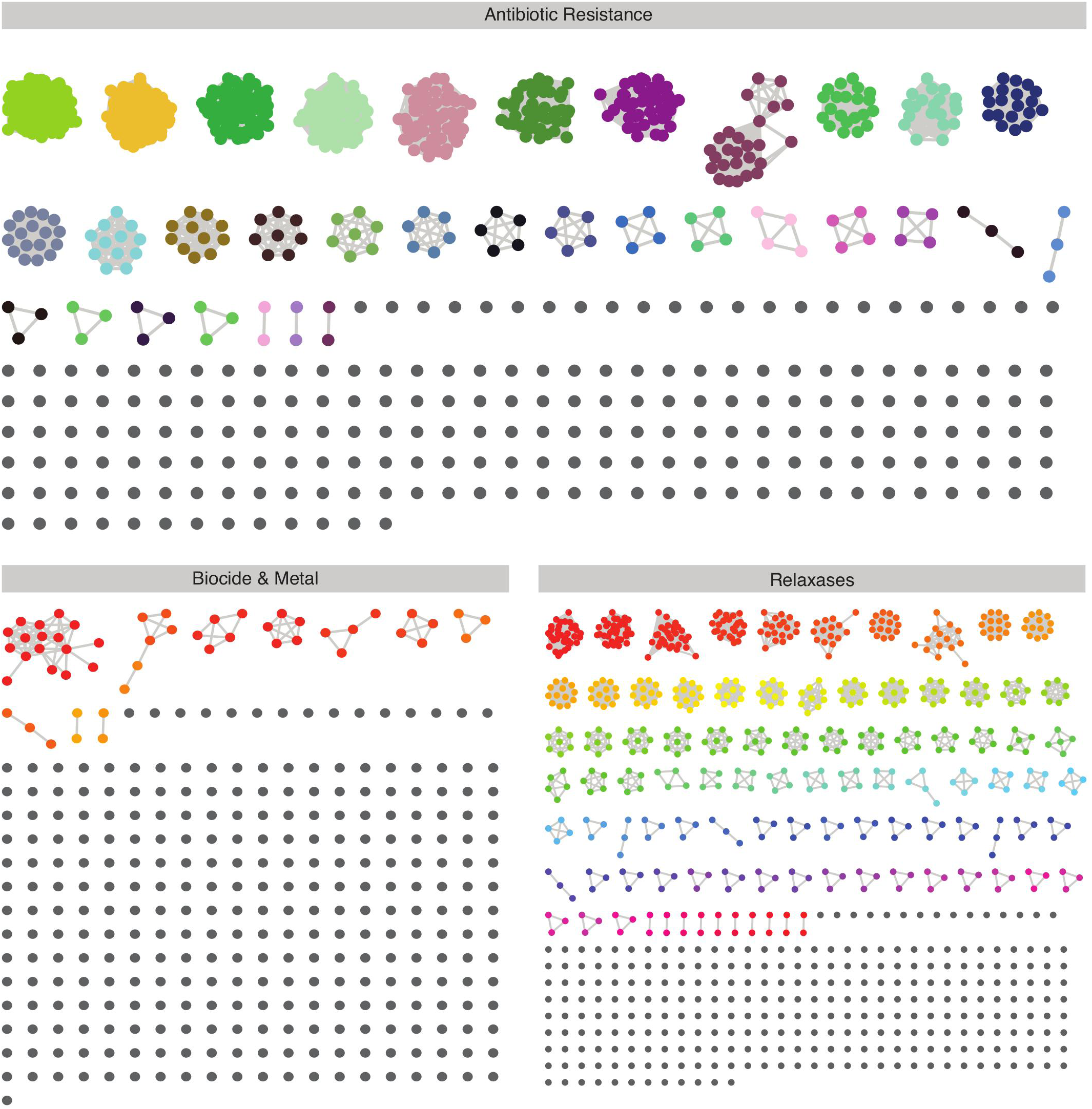
Allele Network: Nodes of the network represents individual genes that are mapped by some read. Edges between nodes represent reads that mapped on both nodes that link. Individual nodes are genes that are unequivocally identified. Gene clusters are mainly composed by different variants of the same gene (alleles). Mapping Gene Cluster (MGC) is defined using Markov cluster algorithm MCL.

